# A gapless Landrace pig genome resolves centromeres and telomeres and highlights telomere repeat structures in different pig breeds

**DOI:** 10.64898/2026.06.25.734473

**Authors:** Harald Grove, Kristina Stenløkk, Sigbjørn Lien, Arne Gjuvsland, Mariann Árnyasi, Maren van Son, Matthew Kent

## Abstract

The Duroc-derived reference genome Sscrofa11.1 has provided a critical foundation for pig genomics, providing a high-quality reference genome for accurate variant detection and comparative genomics but does not capture breed-specific variation. Here, we present a near-complete, gap-free genome assembly for the Landrace pig (*Landrace_v1*, GCA_963921485.1), spanning all 20 chromosomes and totaling 2.6 Gb, including 176 Mb of sequence absent from Sscrofa11.1. Comparative analyses with recently published high-quality pig genomes reveal a conserved centromere organization across breeds, accompanied by substantial variation in repeat composition and length, and identify a pig specific pattern of telomere variant repeats across eight pig breeds. The improved resolution of repetitive regions in *Landrace_v1* enables more complete reconstruction of complex gene families, including olfactory receptors, and uncovers structural variation at the KIT proto-oncogene receptor tyrosine kinase locus not represented in the Duroc reference. Together, these findings highlight the limitations of single-reference genomes and demonstrate the value of breed-specific assemblies for capturing genomic diversity and improving downstream analyses.

## Introduction

Domestic pig (*Sus scrofa*) is a globally important livestock species, serving both as a major source of dietary protein and accounting for ≈30% of global meat consumption (Vicente and Pereira 2024), and as a valuable biomedical model (Lunney, Van Goor et al. 2021). This dual role has motivated extensive genomic research and produced high-quality reference genomes essential for understanding genetic architecture, improving breeding strategies, and enabling translational applications. The first draft reference genome of *S. scrofa* was published in 2012 (Groenen, Archibald et al. 2012) and subsequently improved in 2020 by the Swine Genome Sequencing Consortium who, by incorporating long-read data, resulting in a substantially more contiguous Duroc-derived assembly, Sscrofa11.1 (Warr, Affara et al. 2020). This assembly remains the primary reference for pig genomics. Since then, additional assemblies have been generated for both Asian and European breeds (Zhou, Li et al. 2021, Ma, Jiang et al. 2022, Jiang, Wang et al. 2023), however, many do not yet fully resolve complex genomic regions or key chromosomal features such as centromeres and telomeres. More recently, improved long-read sequencing technologies have enabled a new generation of near-complete, telomere-to-telomere (T2T) genome assemblies (Li and Durbin 2024). High-accuracy HiFi reads from Pacific Biosciences and ultra-long reads from Oxford Nanopore Technologies provide complementary strengths for resolving complex and repetitive regions (Nurk, Koren et al. 2022, Rabanal, Graff et al. 2022, Li and Durbin 2024). These approaches have facilitated the reconstruction of nearly complete chromosomes, including centromeres, telomeres, and other highly variable regions such as immune loci. Several pig breeds have now been assembled to T2T or near-T2T standards, including Min and Rongchang (Zong, Chen et al. 2025); Jinhua (Cao, Miao et al. 2025); Bamei, Juema, Hanjiang Black, Large White (Li, Wang et al. 2024); Wuzhishan (Luo, Huang et al. 2025); Anqing Six-end-white pig (Zhang, Zhou et al. 2025); and the Duroc individual (“Tabasco”) used to build the *S*.*scrofa* 11.1 reference assembly (Li, Wang et al. 2025).

Despite technical advances, closing gaps in genome assemblies remains challenging because these regions are typically enriched for repetitive and structurally complex sequences. Such gaps can disrupt contiguity, lead to misplacement or misorientation of sequences, and obscure biologically important features such as duplicated genes, copy number variants, and regulatory elements (Nurk, Koren et al. 2022, Konig, Mitchell et al. 2024). They also hinder accurate variant detection and genotype imputation, both of which are critical for genomic studies and breeding applications. Centromeres and telomeres exemplify these challenges. Centromeres are typically composed of long arrays of satellite repeats and transposable elements (Kursel and Malik 2016), and are essential for faithful chromosome segregation, and serve as targets in cytogenetic analyses (Donaldson, Villagomez et al. 2021). Complete assemblies are now revealing that centromeric and pericentromeric regions can also contain previously inaccessible structural variation, repeat diversity and gene annotations (Altemose, Logsdon et al. 2022, Pineda, MacPhillamy et al. 2025). Telomeres protect chromosome ends and play key roles in genome stability, aging, and cellular senescence (Whittemore, Vera et al. 2019), with their length and structure influenced by biological and environmental factors (Zhang, Baker et al. 2023). In addition to these chromosome-scale architectural elements, highly duplicated gene families, such as immune genes and olfactory receptor (OR) genes, remain difficult to assemble accurately due to their repetitive and clustered organization, yet are critical for understanding gene function, evolution, and phenotypic variation.

An additional limitation of current genomic resources is that the primary reference genome is derived from a single Duroc individual. While Duroc pigs are widely used as sire lines, other breeds such as Landrace and Large White are commonly used as dam lines and exhibit distinct phenotypic and genomic characteristics (Zhang, Zhang et al. 2020). For example, the dominant white coat color in Landrace pigs is associated with duplication of the KIT proto-oncogene receptor tyrosine kinase (*KIT*) gene (Rubin, Megens et al. 2012), a feature not represented in the Duroc reference. For breed-specific variant detection, reliance on a reference genome from a different breed can introduce reference bias, whereby sequences similar to the reference are more readily detected than divergent or absent sequences (Crysnanto and Pausch 2020). This bias can reduce the accuracy of variant discovery, particularly for structural variants, and affect downstream analyses in breeding programs. These limitations have motivated a shift from single linear reference genomes toward pangenome frameworks that integrate multiple high-quality assemblies, enabling more accurate representation of structural variation, improved read mapping across diverse populations, and enhanced detection of variants in regions that are absent or misrepresented in a single reference.

In this study, we generated a near-complete, gapless reference genome for a Norwegian Landrace pig by integrating long-read sequencing data from Pacific Biosciences and Oxford Nanopore Technologies. The resulting assembly (Landrace_v1, GCA_963921485.1) includes 14 telomere-to-telomere chromosomes, with only minor gaps remaining in the telomeric regions of the others. This assembly resolves complex genomic regions, including centromeres and telomeres, and enables more accurate placement of genes absent from the Duroc reference. Beyond improving assembly completeness, Landrace_v1 provides a more representative genomic framework for white pig breeds, reducing reference bias and enhancing the accuracy of variant detection and structural analysis. Comparative analyses with other high-quality pig genomes further reveal conserved and lineage-specific features of chromosome architecture, including a previously unreported repeat pattern in subtelomeric regions. Together, these advances provide a stronger foundation for genetic studies, trait mapping, and genomic selection in pigs.

## Materials and Methods

### Samples

DNA was extracted from 200 µl of blood collected from a single healthy male Landrace pig at the Norsvin SA breeding center in Hamar, Norway, using the Nanobind CBB Big DNA Kit (Pacific Biosciences). DNA purity was quantified using a NanoDrop 8000 Spectrophotometer, its concentration was measured using Qubit fluorimetry (Thermo Fisher Scientific), and its integrity was subjectively assessed using agarose gel electrophoresis. Fragments <25 kb were progressively depleted using a Short Read Elimination kit (Circulomics), and library preparation was performed using the Oxford Nanopore Technologies (ONT) Ligation Sequencing gDNA Kit (SQK-LSK110) following manufacturers’ instructions. The resulting library was sequenced using two R9.4.1 flow cells on a PromethION-24 sequencer from ONT. From the same individual an additional 5M PacBio HiFi-reads (avg length 16.65Kb; total 82.5Gb) were generated along with 330M short Illumina reads (PE150) by a commercial provider (Novogene UK).

### Data processing and assembly

The ONT reads were converted from fast5 to fastq format using Guppy version 5.1.13 and pre-processed by Filtlong v0.2.1 (https://github.com/rrwick/Filtlong) to remove reads < 4000bp, as well as the 10% of remaining reads with lowest average Q-scores. HiFi-reads were preprocessed with HiFiAdapterFilt v2.0.1 (Sim, Corpuz et al. 2022) using default settings.

Filtered ONT reads were assembled into contigs using Flye (v 2.9.1-b1780) (Kolmogorov, Yuan et al. 2019) with the “--asm-coverage” flag set to 40 to reduce the memory consumption in the initial disjoining step; a range of required read overlaps (--min-overlap flag) were used (5k, 7k, 10k, 15k, 20k, 25k and 30k) producing multiple independent assemblies. Separately, an assembly was created with HiFiasm v 0.19.6-r595 (Cheng, Concepcion et al. 2021, Cheng, Jarvis et al. 2022) using both datasets (filtered HiFi and ONT) and default settings. The resulting HiFiasm contigs were manually connected using overlapping contigs generated by Flye to bridge gaps and generate chromosomes. This was done by aligning the HiFiasm assembly to each Flye assembly and identifying all instances of an overlap between a Flye contig and two HiFi contigs. Candidate joins were accepted only when a Flye contig overlapped both adjacent HiFiasm contigs by at least 10 kb, showed consistent orientation, and was supported by long-read alignments spanning the inferred junction. Any missing sequence between the two HiFi contigs was filled in using the sequence from the Flye contig

The resulting gap-filled assembly was subjected to completeness analysis using Compleasm v0.12-r237 (Huang and Li 2023) with the mammalia_odb10 database containing 9171 expected BUSCO genes which provides values comparable to BUSCO c-scores. Further quality metrics were calculated with QUAST v5.0.2 (Gurevich, Saveliev et al. 2013).

### Synteny

To visualize the differences between assemblies, we mapped Sscrofa11.1 chromosome sequences to Landrace_v1 with minimap2 v2.18 (Li 2018) using the -ax asm5 -eqx flags for whole genome alignments as recommended by Syri v. 1.6.3 (Goel, Sun et al. 2019) which was used to compute synteny; Syri output was plotted using plotsr v1.1.1 (Goel and Schneeberger 2022).

### Localizing putative centromere regions

Searching the NCBI database for satellite DNA and centromere sequence in pigs revealed 25 accessions described as being centromere specific (Supplementary Table 1). The provided annotation of these sequences grouped them into different categories: (i) metacentric repeats (Mc1 and Mc2; 15 sequences) (Jantsch, Hamilton et al. 1990, Riquet, Mulsant et al. 1996), (ii) acrocentric repeats (Ac and Ac2; 6 sequences) (Jantsch, Hamilton et al. 1990) and (iii) unspecified centromeric repeats (centDNA; 4 sequences) (Miller, Hindkjaer et al. 1993). Centromere sequences were mapped to the reference genome using Lastz with default parameters. Alignments with ≥ 90% sequence coverage and ≥ 60% sequence identity were considered probable hits and the fraction of bases covered by one or more query sequence within a 10kb sliding window was calculated across the genome. Windows with more than 10% of the sequence covered by these queries were considered to be a potential part of a centromere region. To supplement this strategy, we catalogued tandem repeats in the genome with Tandem Repeat Finder (TRF, v4.09) software (Benson 1999). Based on the expectation that centromeres will contain an elevated number of repeats, we split the assembly into windows of 1Mbp and calculated the proportion of bases within each window classified as a tandem repeat by TRF.

**Table 1.** Comparison of genome assembly statistics for Landrace_v1 (GCA_963921485.1) and Sscrofa11.1.

|  | **Sscrofa11.1** | **Landrace_v1** |
| --- | --- | --- |
| Accession number | GCA_000003025.6 | GCA_963921485.1 |
| Total genome length | 2,501,895,775 | 2,633,804,377 |
| Sequence in chromosomes | 2,435,262,063 | 2,599,453,217 |
| Gap number | 93 | 0 |
| Contig number | 1117 | 54 |
| Avg contigs per autosomal chromosome; (StdDev) | 5.27 (3.56) | 1 (0) |
| GC (%) | 42 | 42 |
| Compleasm (%) | 99.40 | 99.56 |
| Scaffold N50 | 88,231,837 | 142,898,213 |
| Contig N50 | 48,231,277 | 142,898,213 |
| Scaffold L50 | 9 | 7 |
| Contig L50 | 15 | 7 |

### Localizing telomere regions

Vertebrate telomeres are widely accepted to consist of a variable number of repeating units of the hexamer TTAGGG (Moyzis, Buckingham et al. 1988, Greider 1996). Our approach to identify putative telomeres was to count the number of TTAGGG-repeats within windows of 10kb along the genome. TRF output was also examined for tandem repeats clustering at the ends of chromosomes. We also looked at the first and last 50kb of each chromosome and examined all sequences contained within the telomere repeat-blocks.

### KIT gene investigation

The KIT gene is duplicated in white pigs including Landrace (Rubin, Megens et al. 2012). A whole genome alignment of Landrace_v1 against itself was done using Minimap v2.0 to confirm the duplication. A more detailed look at the KIT region, describing several tandem duplications within the region has been reported (Wu, Deng et al. 2019). They also report a set of primers to identify and count the number of units within each duplication (Supplementary Table 2), and these primers were downloaded and mapped to Landrace_v1 using Last (Kielbasa, Wan et al. 2011) to see if we could determine the exact alleles of the KIT region.

**Table 2.** Mapping of genes annotated in Sscrofa11.1 to Landrace_v1 using the LiftOff genome annotation tool.

|  | **Named** | **LOC** | **orf** | **Total** |
| --- | --- | --- | --- | --- |
|  | 17,419 | 9848 | 339 | 27304 |
| **Contextual match**<br>(ChrA, pos 1 -> ChrA, pos 1) | 16,706 | 8263 | 330 | 25336 |
| **Intrachromosomal change**<br>(ChrA, pos1 -> ChrA, pos2) | 63 | 274 | 2 | 339 |
| **Interchromosomal change**<br>(ChrA -> ChrB) | 14 | 51 | 0 | 65 |
| **New position**<br>(Unmapped contig -> ChrA) | 256 | 978 | 7 | 1241 |
| **Lost from chromosome**<br>(ChrA -> Missing) | 36 | 120 | 0 | 156 |
| **Lost from unmapped**<br>(Unmapped contig -> Missing) | 5 | 162 | 0 | 167 |

### Annotation

Gene annotations from the SusScrofa11.1 reference (version 106, 2017) were downloaded from NCBI and transferred to the Landrace_v1 assembly using LiftOff software (v1.6.3) (Shumate and Salzberg 2021) with default values for alignment coverage (>0.5) and identity (>0.5). Gene ontology enrichment analysis was performed with the PANTHER Overrepresentation Test, using the FISHER test type and keeping the FDR adjusted p-values as the final results. Automated annotation was also performed by EMBL EBI and shared with us for analysis in this paper. This will be made public in Ensembl Release 116.

### Repeat annotation

Repeats in the genome were detected and categorized by RepeatMasker v4.1.1 (Smit, Hubley et al. 2020) using the provided “sus scrofa” database (Storer, Hubley et al. 2021).

### Pan genomic telomere and centromere sequence comparison

We downloaded seven pig assemblies representing a diversity of global breeds where each is reported to be of telomere-to-telomere quality or close to it (Supplementary Table 3). To enable comparative analysis, centromeres and telomeres for each assembly were detected using the methods described above. The goal was not to try and describe the centromere and telomeres in the other breeds but rather have a measure to compare how similar these other assemblies are to the new Landrace assembly. T2T assemblies for cattle, human and sheep served as cross-species comparators (Supplementary Table 3).

## Results

### Hybrid assembly strategy yields a near-complete, gapless Landrace pig genome

To construct the first high-quality genome assembly of the Norwegian Landrace pig, we generated long-read sequencing data from an individual male using Oxford Nanopore Technologies (ONT; 123.5 Gb, ~45× coverage; read N50 = 42.7 kb) and Pacific Biosciences (HiFi; 87.5 Gb, ~35× coverage; read N50 = 16.2 kb). Reads were quality filtered and used to generate two preliminary assemblies with Hifiasm and Flye. These were merged to create a near-complete, gapless genome comprising 2.6 Gb of chromosome sequence. To assess whether short-read polishing could further improve assembly quality, we generated and incorporated 330 million Illumina paired-end reads. However, incorporating these data produced no measurable improvement in standard assembly-quality metrics. The final assembly, Landrace_v1 (GCA_963921485.1; Table 1), therefore represents a chromosome-scale genome reconstruction based exclusively on long-read sequencing data.

### Extensive novel sequence and structural variation revealed by comparison to the Duroc reference genome

Comparing the length of the Landrace assembly to the widely used Duroc reference genome (Sscrofa11.1) revealed the additions of 176 Mb of new sequence and the loss of 12 Mb giving a net change of 164 Mb (6.2%) of chromosome-anchored sequence. Synteny analysis (Figure 1) indicates that the majority of this sequence co-localizes with, extends, or fully bridges previously unresolved centromeric regions in metacentric chromosomes Ssc01–Ssc11 and telo-/acrocentric chromosomes Ssc13, 15, 17, and 18. Moreover, centromeric regions are now resolved in Ssc12 and Ssc14, and partially resolved in Ssc16, where corresponding sequence was absent or not annotated in Sscrofa11.1.

**Figure 1.**
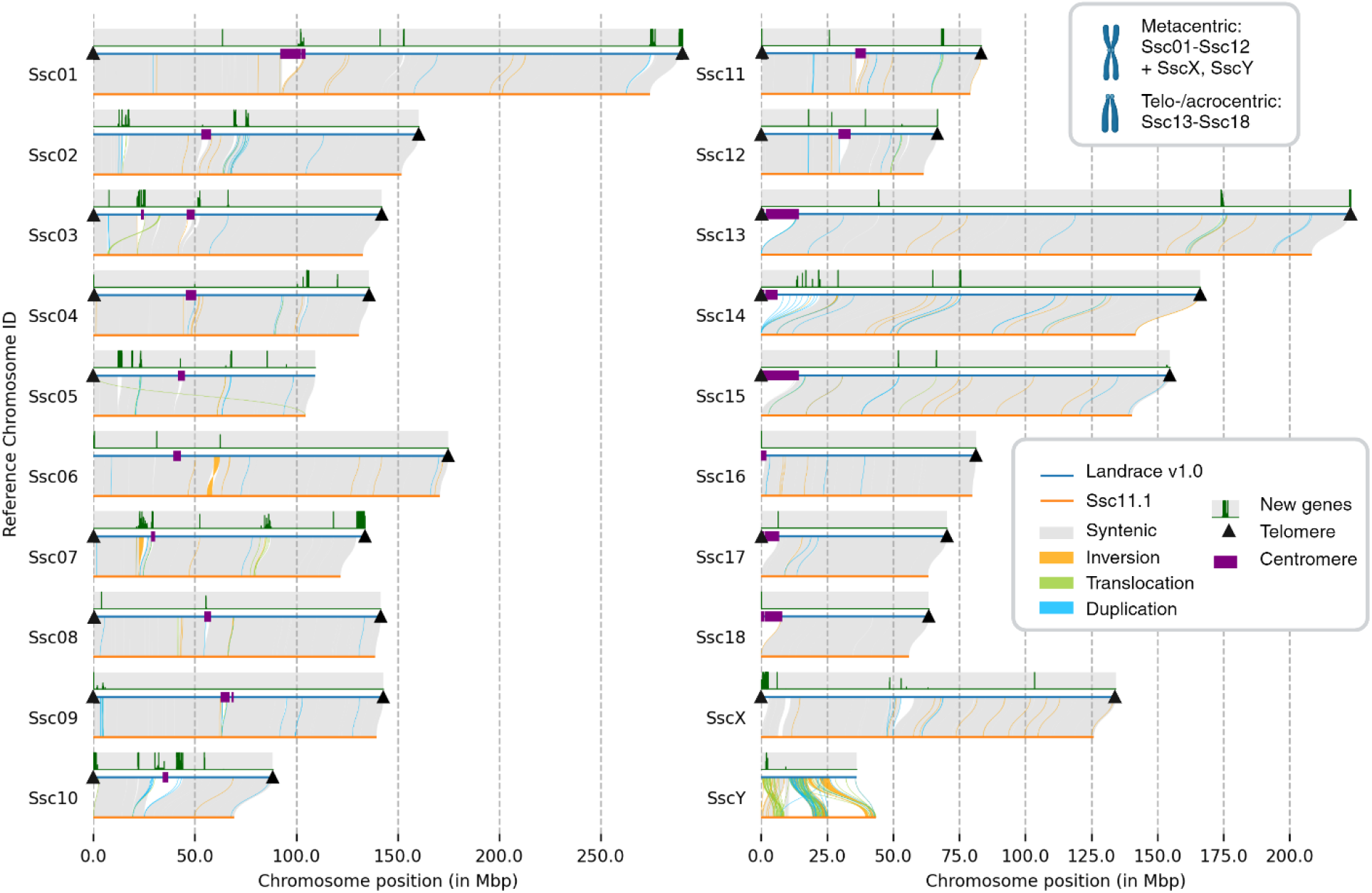
Synteny between Landrace_v1 and Sscrofa11.1. Predicted telomeres (triangles) and centromeres (purple blocks) are shown, as visualized using the Python package plotsr. Genes relocated from previously unmapped contigs in Sscrofa11.1 to chromosomal positions in Landrace_v1 are indicated in dark green. Chromosomes Ssc01–Ssc12, SscX, and SscY are metacentric, whereas Ssc13–Ssc18 are telo-/acrocentric.

A more detailed analysis of whole-genome alignments identified 176 Mb of novel sequence distributed across 877 regions (Supplementary Table 4), indicating that additional sequence is also present outside canonical centromeric regions. The amount of new sequence per chromosome ranged from 0.74 to 23.5 Mb, with chromosome 10 exhibiting the largest relative increase in size (20%). Structural variation analysis using SyRI structural variant tool further revealed extensive genomic rearrangements, including 230 inversions, 153 translocations, and 629 duplications. These variants collectively affect 17 Mb, 18 Mb, and 6 Mb of sequence, respectively, highlighting substantial structural divergence between the Landrace assembly and the Sscrofa11.1 reference.

### Improved genome completeness enables gene reassignment and refined annotation in Landrace_v1

*De novo* annotation of the Landrace_v1 assembly identified 19,252 genes and 440 pseudogenes. Overall, gene content was highly similar to that of the Sscrofa11.1 reference, with the main differences being an increase in lncRNA and ncRNA gene features, as well as in C-, D-, J-, and V-segment annotations associated with immune function (Supplementary Table 5). To enable direct comparison between assemblies, we performed gene liftover to transfer Sscrofa11.1 annotations onto Landrace_v1. This analysis reassigned 1,241 genes from previously unmapped contigs in Sscrofa11.1 to chromosomal positions in Landrace_v1, reflecting improved genome completeness. Most of these genes (84%) are uncharacterized LOC genes lacking assigned function, whereas 256 correspond to annotated genes. An additional 339 genes were repositioned within chromosomes and 65 relocated between chromosomes, while 323 genes were not detected in the Landrace_v1 assembly (Table 2).

To assess the functional relevance of newly placed genes (“New position” in Table 2), we performed gene ontology enrichment analysis (Supplementary Table 6), which revealed significant enrichment for olfactory-related processes. This finding is consistent with the known expansion and high sequence similarity of olfactory receptor gene families in mammalian genomes, which are often difficult to assemble due to their repetitive nature (Barnes, Ibarra-Soria et al. 2020). In contrast, genes absent from Landrace_v1 showed no clear pattern of regional clustering or functional enrichment.

We further examined gene content within sequence present in landrace_v1 but absent from Sscrofa11.1 by intersecting regions identified by SyRI structural variant tool with Ensembl annotations. Olfactory receptor (OR) genes represented the largest gene class in these regions: although they constitute approximately 3% of all genes, they account for 21% of genes located within newly identified sequences. This enrichment is particularly pronounced on chromosome Ssc07, which contains 31% of newly identified OR genes despite harboring only 18% of the total OR gene repertoire.

### Conserved centromere organization with breed-specific variation in repeat composition and structure

To assess the completeness and structural organization of centromeres in the *Landrace_v1* assembly, we mapped canonical centromere sequences (Supplementary Table 1) to the genome and identified regions enriched for centromeric content using 10 kb windows (>10% centromeric sequence). Most chromosomes contained a single enriched region, consistent with expected centromere organization, whereas chromosomes Ssc03 and Ssc06 each exhibited two such regions (Supplementary Figure 1). The canonical sequences mapped to multiple genomic locations with sequence similarities ranging from 60% to 90%, indicating substantial divergence between reference centromeric sequences and those present in the Landrace genome.

To benchmark these assignments, we compared inferred centromere positions and lengths (Supplementary Table 7) with published data from two Chinese pig breeds, Min and RC (Zong, Chen et al. 2025). Centromere positions were highly concordant across datasets (r^2^ > 0.99), whereas centromere lengths showed greater variability, correlating well with the Min breed (r^2^ = 0.81) but less so with the RC breed (r^2^ = 0.28), suggesting breed-specific differences in centromere expansion, repeat composition or assembly differences.

Analysis of centromere-associated repeat families revealed distinct patterns between chromosome types. Acrocentric-specific repeats (Ac2), with unit lengths ranging from 76 to 357 bp, mapped exclusively to the proximal regions of acrocentric chromosomes (Ssc13–Ssc18). In contrast, metacentric repeats (Mc1: 190–313 bp; Mc2: 191–565 bp) were enriched in metacentric chromosomes, with Mc2 repeats predominantly localized to Ssc04–Ssc11 and co-occurring with Mc1 repeats, which were detected across most chromosomes except Ssc16. Three previously uncategorized ~336 bp repeat sequences co-localized with Mc1 repeats and exhibited the highest sequence identity to the assembly; these were therefore grouped within the Mc1 class for subsequent analyses. Among the remaining Mc1 sequences, only a subset (X51555.1, X51559, and X51560.1) showed consistent mapping. The distribution of these repeat families is consistent with the expected karyotype in which chromosomes Ssc01–Ssc12, SscX, and SscY are metacentric, whereas Ssc13–Ssc18 are acrocentric.

Regions enriched for centromeric sequence showed strong concordance with regions containing a high density of tandem repeats. Using Tandem Repeats Finder, we quantified tandem repeat (TR) content in 1 Mb windows (Supplementary Figure 1). These regions were dominated by two principal repeat types: ~336 bp units and ~50 bp units, co-localizing with the Mc1 satellites and the Mc2 satellites respectively. The 50 bp repeats displayed substantial sequence variation both between and within chromosomes, which in some cases prevented consistent classification into a single repeat family.

Together, the centromere-specific repeat mapping and tandem repeat analyses indicate that metacentric centromeres in the Landrace genome are composed either of homogeneous arrays of Mc1 (~336 bp) repeats or of mixed arrays containing both Mc1 (~336 bp) and Mc2 (~50 bp) repeats (Figure 2).

**Figure 2.**
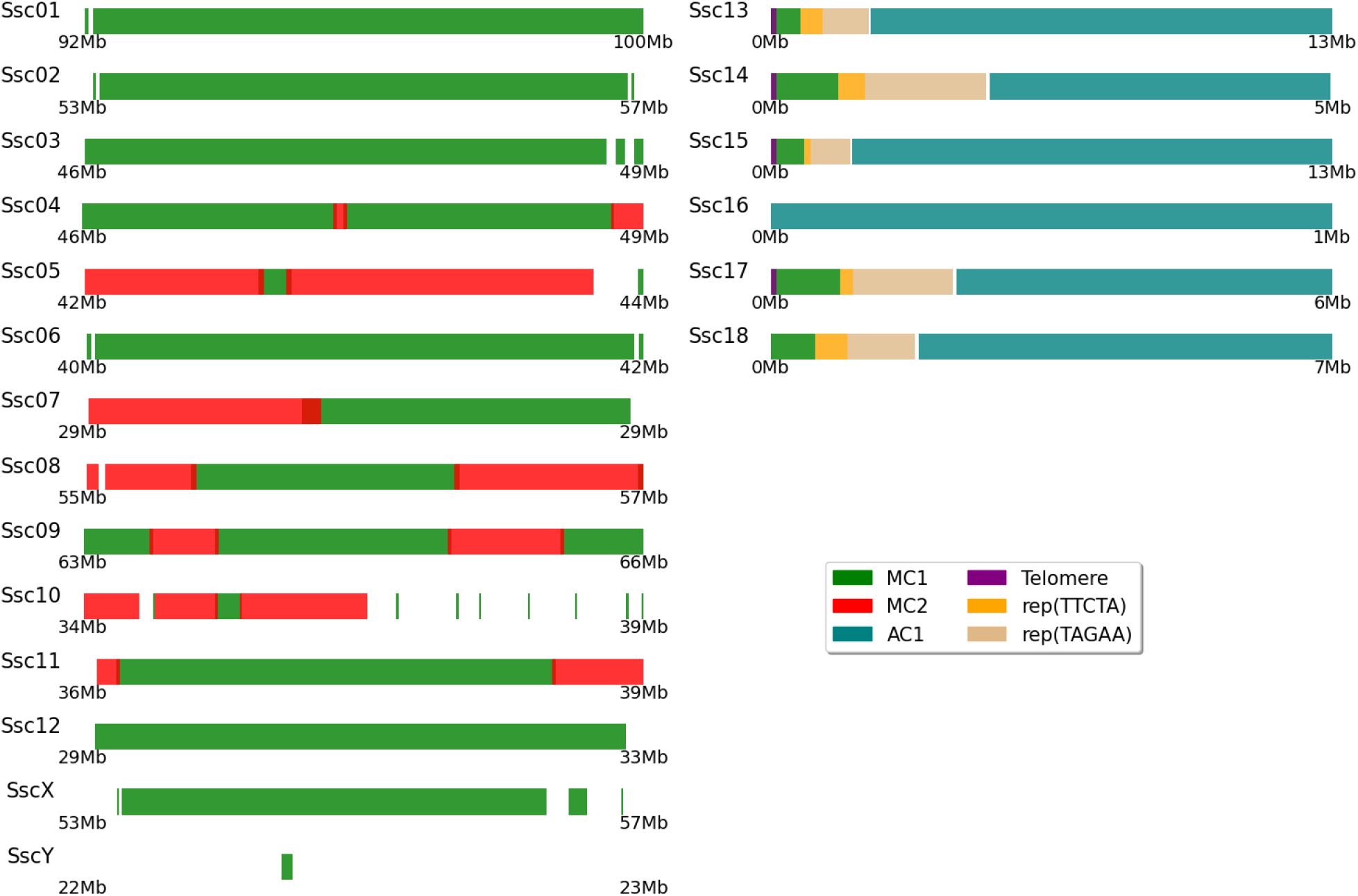
Schematic representation of the inferred centromere structure based on the distribution of MC1, MC2, and AC2 repeat families. The model also illustrates the organization of tandem repeats at the proximal ends of acrocentric chromosomes (Ssc13–Ssc18), excluding Ssc16.

The proximal ends of all acrocentric chromosomes (Ssc13–Ssc18), with the exception of Ssc16, where sequence does not extend fully to the chromosome start, show a consistent structural organization. Each begins with a telomeric repeat tract, followed by an Mc1-enriched region, a tandem repeat block containing the pentamer TTCTA, a corresponding block of its reverse complement (TAGAA), and finally the Ac2 satellite array (Supplementary Table 8).

Applying this centromere classification framework to seven additional pig T2T genomes revealed a broadly conserved organization across breeds, with the primary variation arising from differences in the length of individual repeat segments (Figure 3 and Supplementary Figure 2).

**Figure 3.**
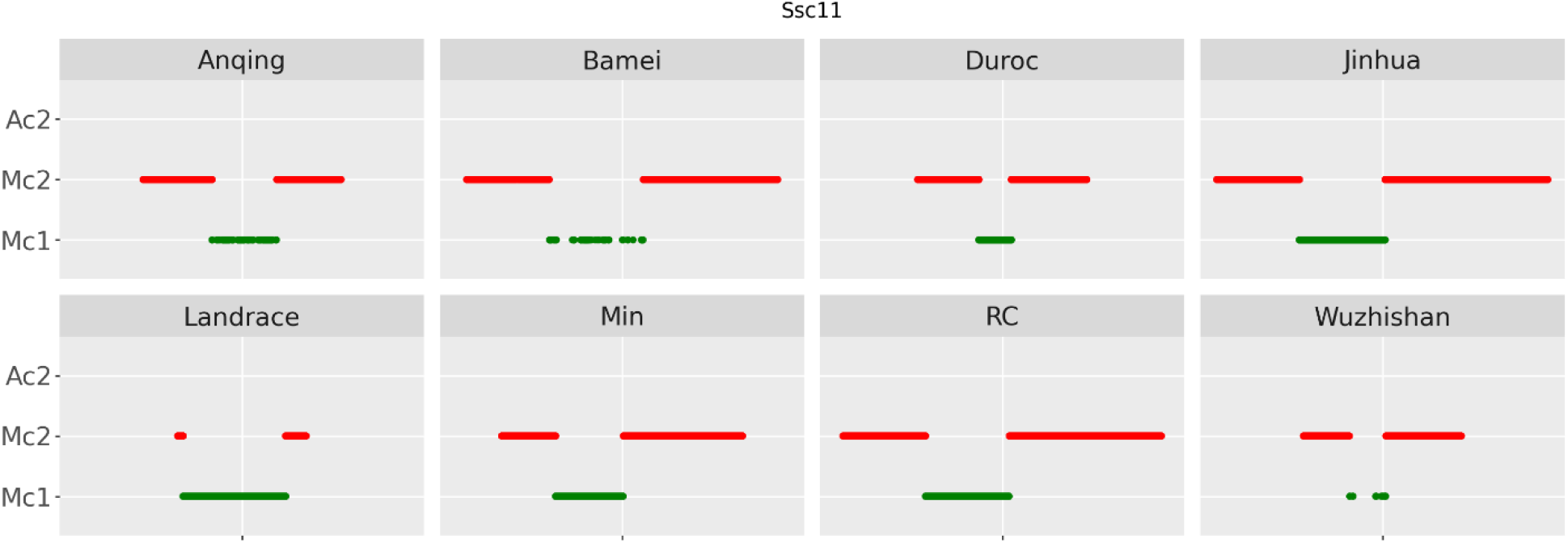
Composition of the centromere on chromosome Ssc11 across Landrace_v1 and seven additional pig genomes. Most breeds show a conserved centromere position and repeat organization, characterized by an Mc2/Mc1/Mc2 pattern, although the lengths of the individual repeat segments vary between breeds.

Notably, the centromere of chromosome Ssc03 in the Anqing pig is located within an inverted region relative to the corresponding region in other breeds, highlighting a distinct structural rearrangement.

### Conserved subtelomeric architecture reveals porcine telomere-associated repeat motifs

The canonical telomeric repeat (TTAGGG) was detected at both ends of most chromosomes in the Landrace_v1 assembly (Figure 1), consistent with expected telomere structure. However, at the transition between telomeric repeats and adjacent chromosomal sequence, we identified additional short repeat motifs of 5–6 bp that closely resemble the canonical telomere sequence (Figure 4).

**Figure 4.**
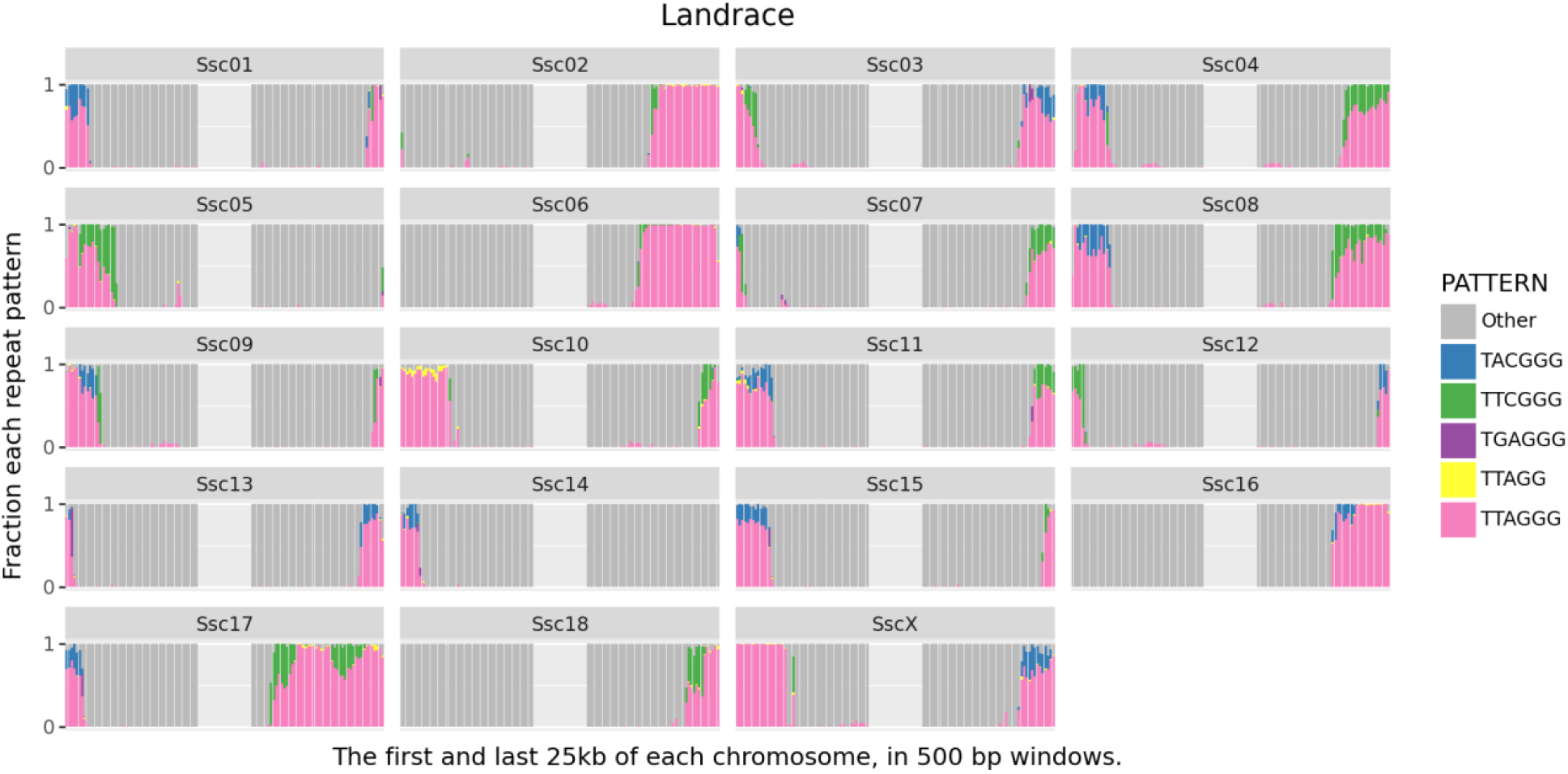
Telomeric regions in Landrace_v1. The first and last 25 kb of each chromosome are shown, divided into 500 bp windows. Each bar represents the proportion of repeat types within a window. Gray indicates sequence not assigned to known repeat classes, while pink denotes canonical telomeric repeats. Most chromosomes also contain additional repeat motifs interspersed with the telomeric sequence.

These resemble the Telomere Variant Repeats (TVR) found between the telomere and the sub-telomere in human genomes (Conomos, Stutz et al. 2012, Sieverling, Hong et al. 2020, Stephens and Kocher 2024). These TVRs were typically dominated by a single repeat per telomere, although their abundance and block length varied. This pattern was consistently observed in Landrace_v1 as well as across seven additional pig genomes (Figure 5), indicating that it represents a conserved feature of porcine telomere architecture.

**Figure 5.**
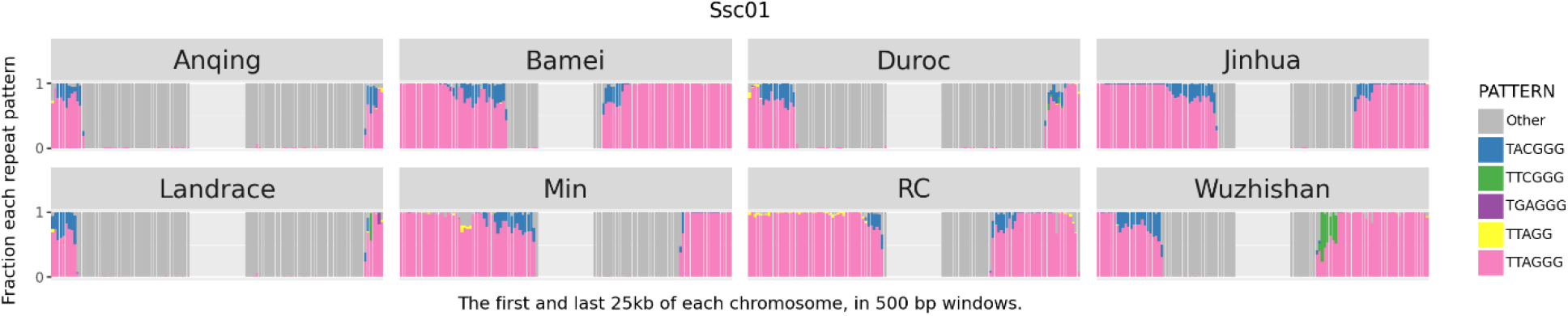
Telomeric region of chromosome Ssc01 across eight pig breeds. The first and last 25 kb of each chromosome are shown, divided into 500 bp windows. Each bar represents the proportion of repeat types within a window. Gray indicates sequence not assigned to known repeat classes, while pink denotes canonical telomeric repeats. Most breeds display a TACGGG repeat at the transition between the telomeric repeat (TTAGGG) and adjacent sequence. The length of the telomeric repeat tract varies between breeds. Corresponding plots for additional chromosomes are provided in Supplementary Figure 3.

While the predominant motif in subtelomeric regions was generally conserved across genomes, some variation was evident, suggesting limited sequence divergence between breeds in these regions. In addition to terminal locations, the telomeric repeat (TTAGGG) was also enriched at internal positions on chromosomes Ssc01, Ssc03, Ssc06, Ssc07, and Ssc08.

### The KIT locus reveals breed-specific structural variation in Landrace

To evaluate the representation of a known breed-specific locus, we examined the assembly of the KIT proto-oncogene receptor tyrosine kinase (*KIT*) gene, which underlies the dominant white phenotype in Landrace pigs. This locus is known to be duplicated in the Landrace genome (Rubin, Megens et al. 2012), whereas it is present as a single copy in the Duroc-derived Sscrofa11.1 reference (Johansson Moller, Chaudhary et al. 1996).

Inspection of the corresponding region in the whole-genome alignment between Landrace_v1 and Sscrofa11.1 confirmed the expected duplication spanning the *KIT* locus (Supplementary Figure 4). Previous studies have described multiple alleles at this locus, defined by variation in the copy number of three smaller CNVs embedded within the larger duplicated region. To characterize this structure in Landrace_v1, we used published primer sequences (Supplementary Table 2) to identify the corresponding breakpoints and infer the copy numbers of the different CNVs. While the overall structural organization of the locus was consistent with previously reported alleles, differences were observed in the number of copies of the individual CNV segments (Figure 6), indicating that the Landrace_v1 assembly represents a distinct allele at the *KIT* locus.

**Figure 6.**
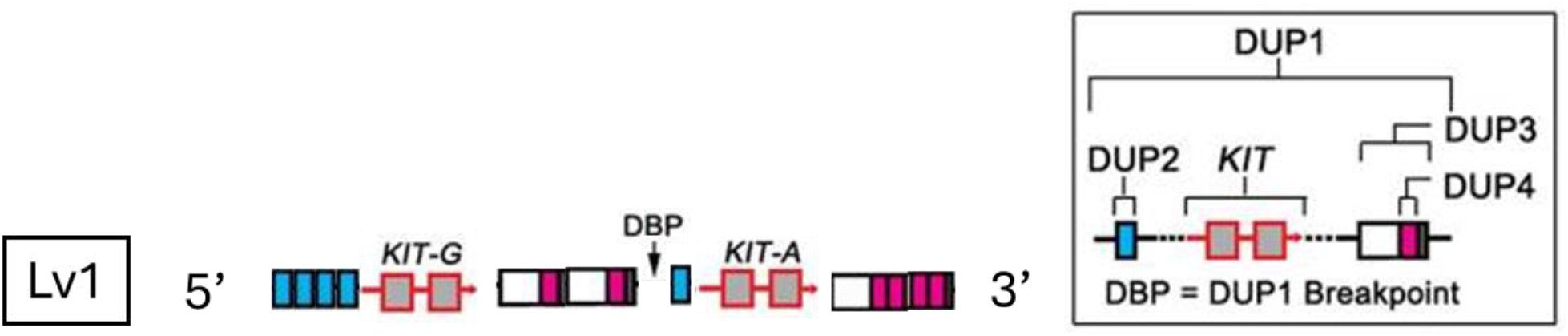
*In silico* characterization of the KIT proto-oncogene receptor tyrosine kinase locus using primer sets reported by Rubin et al. and Wu et al. (Rubin, Megens et al. 2012, Wu, Deng et al. 2019). Symbol annotations follow the conventions defined in these studies. While the overall structural organization of the locus is consistent with previously described alleles, the copy number of the DUP2, DUP3, and DUP4 segments differs, indicating the presence of a novel allele in Landrace_v1.

### Repeat composition of Landrace_v1 is broadly conserved relative to Sscrofa11.1

A total of 28.62% (743.9 Mb) of the Landrace_v1 chromosome sequence was annotated as repetitive. The most abundant class comprised long interspersed nuclear elements (LINEs), accounting for 17.42% of the genome, with the LINE1 family representing 14.33%. Long terminal repeat (LTR) elements were the second most prevalent class, covering 3.5% of the genome. The remaining repeat categories, including SINEs, DNA transposons, small RNA-associated repeats, simple repeats, and low-complexity or unclassified sequences, collectively accounted for 7.69% of the genome. Comparison with the Sscrofa11.1 reference indicated that overall repeat composition is largely conserved, with no major differences observed between the assemblies.

## Discussion

We present a near-complete, gapless, breed-specific reference genome for the Landrace pig (Landrace_v1), generated using two complementary long-read sequencing technologies. Compared to the widely used Duroc-derived reference (Sscrofa11.1), this assembly captures substantial additional sequence and structural complexity, underscoring the limitations of relying on a single linear reference genome for a genetically diverse species. Our results align with a growing consensus in genomics that population- or breed-specific assemblies are essential for reducing reference bias and improving the accuracy of variant discovery, comparative analyses, and downstream applications (Li, Wang et al. 2025, Ren 2026).

A central advance of Landrace_v1 is the resolution and detailed characterization of centromeric regions across all chromosomes. By integrating canonical centromere sequence mapping with tandem repeat analysis, we show that centromeres are consistently positioned but vary markedly in size and repeat composition. Metacentric chromosomes are dominated by Mc1 or mixed Mc1/Mc2 arrays, whereas acrocentric chromosomes exhibit a distinct Ac2-associated structure. Comparative analyses with other pig T2T assemblies demonstrate that this organizational framework is conserved across breeds, despite substantial variation in repeat length and sequence composition. These findings indicate that centromere architecture is evolutionarily stable at the structural level but dynamic at the sequence level. This duality has important implications for chromosome evolution, as variation in centromere structure can shape recombination landscapes and contribute to the formation of structural variants (Ducos, Berland et al. 2007), including inversions identified in this study.

Telomere analyses further revealed previously uncharacterized features at chromosome ends. In addition to canonical telomeric repeats, we identified conserved short repeat motifs in sub-telomeric regions that are shared across multiple pig genomes. This phenomenon has been described as telomere variant repeat (TVR) regions in humans and is found between the telomeres and the sub-telomeres (Conomos, Stutz et al. 2012, Sieverling, Hong et al. 2020, Stephens and Kocher 2024). TVRs are more common in cells with alternative lengthening of telomeres (ALT) (Conomos, Stutz et al. 2012, Lee, Hills et al. 2014) and do not bind the proteins needed for the telomerase operation, instead promoting telomere-telomere recombination (Marzec, Armenise et al. 2015). The eight pig genomes (including Landrace_v1) had predominantly two TVRs: TTCGGG and TACGGG. Both have been reported for humans but the second one with low frequency (Sieverling, Hong et al. 2020). We did not observe the TVR region in the genomes from cattle and sheep, but we also found just on TVR (TGAGGG) in three of the telomeres from the human genome we investigated. There may be an effect of which cell type that was used for DNA extraction, but further work will be required to answer the question of why we observed this pattern only in the pig genomes. We also observed substantial variation in telomere length across chromosomes, highlighting the dynamic nature of these regions. This variability is likely influenced by biological and environmental factors, such as age (Zhang, Baker et al. 2023), tissue type (Jiang, Carter et al. 2004), and physiological stress (Dugdale and Richardson 2018), although genetic effects may also contribute. While this complicates direct functional interpretation, it underscores the need for further investigation into the role of telomere dynamics in genome stability and phenotypic variation.

Beyond chromosome-end and centromere structure, the improved assembly substantially enhances the representation of complex gene families. In particular, the enrichment of olfactory receptor (OR) genes within newly resolved sequence highlights the value of long-read sequencing for capturing highly duplicated and repetitive loci that are often fragmented or collapsed in earlier assemblies. The OR genes in the pig were reported to include 1113 genes and 188 pseudogenes (Nguyen, Lee et al. 2012). Given the roles of these genes in sensory perception, behavior, and environmental interaction (Rorvang, Schild et al. 2023), their improved representation has clear functional relevance and may enhance the power of genome-wide association studies and other functional genomics approaches.

Analysis of the KIT proto-oncogene receptor tyrosine kinase locus provides a clear example of the value of a breed-specific assembly. The improved resolution afforded by *Landrace_v1* enables detection of allelic diversity consistent with previous reports for Landrace pigs (Rubin, Megens et al. 2012, Wu, Deng et al. 2019), but not captured in the Duroc reference, illustrating how reliance on a single reference can obscure structurally complex variation. Resolving such variation is important not only for traits such as coat color but also for other phenotypes influenced by complex genomic architecture, emphasizing the need for accurate representation of duplicated and structurally complex loci.

Whole-genome alignment of Landrace_v1 to Sscrofa11.1 revealed extensive structural differences, including inversions, translocations, and duplications. While some of these differences likely reflect true biological divergence between Landrace and Duroc breeds, others arise from improved resolution of previously unresolved regions. Regardless, these findings demonstrate that important genomic features, including variants with potential phenotypic effects, can be missed when relying on incomplete or mismatched reference genomes. This has direct implications for the reinterpretation of existing genomic datasets and highlights the importance of continually updating reference resources as sequencing technologies advance.

From an applied perspective, these improvements have important implications for pig breeding and genomics. Modern breeding programs increasingly depend on genomic prediction, variant discovery, and genotype imputation, all of which rely on the quality and representativeness of the reference genome. Using a reference derived from a different breed can introduce bias, reduce mapping accuracy, and impair detection of structurally complex variants. In contrast, a breed-specific assembly such as Landrace_v1 provides a more appropriate framework for identifying causal variants, improving imputation accuracy, and enabling more precise selection strategies.

Taken together, our findings support a transition from single-reference genomes toward pangenome frameworks that integrate multiple high-quality assemblies. Such approaches are increasingly recognized as necessary to capture the full spectrum of genetic diversity within species, including both shared and breed-specific variation (Du, Zhuo et al. 2025, Li, Wang et al. 2025, Ren 2026). The *Landrace_v1* genome represents an important addition to this effort and provides a valuable resource for advancing studies of genome structure, evolution, and breeding in pigs.

## Data availability

The Landrace_v1 reference genome is available with accession number GCA_963921485.1 in the NCBI genome database (https://www.ncbi.nlm.nih.gov/datasets/genome/GCA_963921485.1/).

