## Supplemental Data for "A gapless Landrace pig genome resolves centromeres and telomeres and highlights telomere repeat structures in different pig breeds"

### Supplementary information

#### Supplementary tables

Supplementary Table 1: Entries specified as being centromere related, based on searching NCBI for "satellite DNA" or "centromere sequence repeat", and "Sus Scrofa".

| Accession number | Length | Category | Description |
|---|---|---|---|
| X51555.1 | 313 | Mc1 | S. scrofa Mc1 satellite DNA (clone pAv1.5) |
| X51556.1 | 305 | Mc1 | S. scrofa Mc1 satellite DNA (clone pAv1.6) |
| X51557.1 | 301 | Mc1 | S. scrofa Mc1 satellite DNA (clone pPv15) |
| X51558.1 | 294 | Mc1 | S. scrofa Mc1 satellite DNA (clone pAL7.5) |
| X51559.1 | 190 | Mc1 | S. scrofa Mc1 satellite DNA (clone pAL7.1) |
| X51560.1 | 192 | Mc1 | S. scrofa Mc1 satellite DNA (clone pAL7.4) |
| U42362.1 | 565 | Mc2 | Sus scrofa centromere-specific repeat, T27M clone, Mc2 satellite DNA amplified from S0048 primer set |
| U42363.1 | 549 | Mc2 | Sus scrofa centromere-specific repeat, T27R clone, Mc2 satellite DNA amplified from S0048 primer set |
| U42364.1 | 562 | Mc2 | Sus scrofa centromere-specific repeat, T32M clone, Mc2 satellite DNA amplified from S0048 primer set |
| U42365.1 | 565 | Mc2 | Sus scrofa centromere-specific repeat, T32R clone, Mc2 satellite DNA amplified from S0048 primer set |
| U42366.1 | 277 | Mc2 | Sus scrofa centromere-specific repeat, 22M subclone, Mc2 satellite DNA from S0048 |
| U42367.1 | 277 | Mc2 | Sus scrofa centromere-specific repeat, 29M subclone, Mc2 satellite DNA from S0048 |
| U42368.1 | 197 | Mc2 | Sus scrofa centromere-specific repeat, 29R subclone, Mc2 satellite DNA from S0048 |
| U42369.1 | 351 | Mc2 | Sus scrofa centromere-specific repeat, 30M subclone, Mc2 satellite DNA from S0048 |
| U42370.1 | 191 | Mc2 | Sus scrofa centromere-specific repeat, 30R subclone, Mc2 satellite DNA from S0048 |
| X51561.1 | 309 | Ac2 | S. scrofa Ac2 satellite DNA (clone pMb3.4) |
| X51562.1 | 328 | Ac2 | S. scrofa Ac2 satellite DNA (clone pMb3.5) |
| X51563.1 | 235 | Ac2 | S. scrofa Ac2 satellite DNA (clone pMb2.3) |
| X51564.1 | 156 | Ac2 | S. scrofa Ac2 satellite DNA (clone pHi4.2) |
| X51565.1 | 76 | Ac2 | S. scrofa Ac2 satellite DNA (clone pHi4.5) |
| X62138.1 | 337 | centDNA | S.scrofa centromere-specific repeat |
| X62139.1 | 336 | centDNA | S.scrofa centromere-specific repeat |
| X62140.1 | 335 | centDNA | S.scrofa centromere-specific repeat |
| X70941.1 | 313 | centDNA | S.scrofa DNA for centromere specific motif |
| D11085.1 | 357 | Ac | Sus scrofa domestica DNA, satellite region in centromere of acrocentric chromosomes |

Supplementary Table 2: The selected primer sequences from (Wu et al. 2019) used to categorize the regions of duplications around the KIT gene.

| Name | Primer sequence |
|---|---|
| DUP1_F | GGATGCTACATGCAGCTGTT |
| DUP1_R | GTTTGATGATGTGGGCTGAG |
| DUP1_P | TCAAGCTCACCACCTGGCATTC |
| DUP2_F | TGTGACCAGGAAGGGTAGTG |
| DUP2_R | CAGGTGGGGCTTCTCTCTAT |
| DUP2_P | CCTCACTGCTCCTGCTAATTCCTC |
| DUP3_F | GGACGCATAAATTGTATGTAGATTG |
| DUP3_R | TTCATTTTTGCACATTCCTTCT |
| DUP3_P | TGCCTGCTAACAATTATGCATGCA |
| DUP4_F | GCTCAGGGACCACAGGTATT |
| DUP4_R | TATTTGGCAGGTGTTGGAAA |
| DUP4_P | AGCCTTCTAGCAGCTGATTGGATGG |
| ER_F | ACGGTGGATATGGTCCTTCT |
| ER_R | CGCTCTCTCTCTGCACATTC |
| ER_P | TCAGGGTGCTGGACAGAAACG |
| DUP1BP_F | ATGTGGAGAAGCAGGAGACC |
| DUP1BP_R | TGTTTCACCCGCATCCTACT |
| DUP2BP_F | GACAAGTGTGAAAACAGAAGTGTTA |
| DUP2BP_R | TGTTTACATGCCAGGGGTTCAG |
| DUP3BP_F | AGGCTTGGCACTCAGATGAC |
| DUP3BP_R | TCAGCCTCCCATCACAGGTA |
| DUP4BP_F | CAGTGGTGATGGAGGCTTGG |
| DUP4BP_R | GGGGTTAGGCCTTAAGATGGT |

Supplementary Table 3: Publicly available genomes used for comparison of telomere regions.

| Accession number | Pig Breed | Reference |
|---|---|---|
| GCA_030704935.2 | Bamei | Li et al. 2024 |
| GCA_048338725.1 | Wuzhishan | Luo et al. 2025 |
| GWHFPUM00000000.1 | Rongchang (100 bp gap x 1) | Zong et al. 2025 |
| GWHFPUN00000000.1 | Min | Zong et al. 2025 |
| PRJNA1238047 | Jinhua | Cao et al. 2025 |
| GCA_050231125.1 | Anqing (100 bp gap x 23) | Zhang et al. 2025 |
| GCA_000003025.7 | Duroc | Li et al. 2025 |
| GCF_009914755.1 | T2T-CHM13v2.0 (Human) | |
| GCA_040805955.1 | T2T-sheep1.0 (Sheep) | |
| GCA_040286185.1 | UOA_Wagyu_1 (Bovine) | |
| GCA_051106085.1 | Guyuan_Btau_1.0 (Bovine) | |

Supplementary Table 4: Syri output, showing the total amount of sequence within new, duplicated, inverted and translocated regions. Presented as base pairs per chromosome.

| CHROM | TOTAL [bp] | NEW [bp] | DUP [bp] | INV [bp] | TRANS [bp] |
|---|---|---|---|---|---|
| Ssc01 | 290,485,937 | 15,821,692 | 133,946 | 453,990 | 8,889 |
| Ssc02 | 160,425,991 | 8,645,815 | 363,558 | 315,134 | 19,180 |
| Ssc03 | 142,074,565 | 8,946,410 | 118,591 | 42,650 | 744,116 |
| Ssc04 | 135,905,078 | 1,054,756 | 155,357 | 193,511 | 63,585 |
| Ssc05 | 109,333,974 | 4,548,162 | 252,321 | 83,072 | 223,451 |
| Ssc06 | 174,868,856 | 739,645 | 174,836 | 3,295,659 | 9,945 |
| Ssc07 | 133,832,580 | 9,816,388 | 291,049 | 2,447,878 | 345,858 |
| Ssc08 | 141,541,156 | 2,394,511 | 110,763 | 269,794 | 291,243 |
| Ssc09 | 142,898,213 | 3,088,410 | 449,356 | 71,669 | 83,972 |
| Ssc10 | 88,473,292 | 17,492,082 | 498,424 | 26,394 | 102,881 |
| Ssc11 | 83,339,412 | 4,558,097 | 205,965 | 152,934 | 91,965 |
| Ssc12 | 66,765,734 | 5,295,811 | 192,164 | 45,466 | 149,336 |
| Ssc13 | 223,124,803 | 15,378,505 | 301,940 | 89,923 | 233,436 |
| Ssc14 | 166,181,556 | 23,531,892 | 469,572 | 714,832 | 42,793 |
| Ssc15 | 154,681,081 | 14,010,881 | 228,538 | 268,940 | 15,620 |
| Ssc16 | 81,336,253 | 1,671,195 | 137,969 | 67,917 | 8,411 |
| Ssc17 | 70,321,686 | 6,740,313 | 102,407 | 104,595 | 8,255 |
| Ssc18 | 63,460,842 | 7,595,088 | 35,086 | 13,145 | 13,678 |
| SscX | 134,215,034 | 7,633,630 | 1,273,291 | 485,106 | 7,126,003 |
| SscY | 36,170,416 | 16,929,928 | 758,976 | 7,841,061 | 8,886,344 |
| SUM | 2,599,436,459 | 175,893,211 | 6,254,109 | 16,983,670 | 18,468,961 |

Supplementary Table 5: List of features in the new annotation of Landrace_v1, compared to the annotation from Sscrofa11.1 (both Ensembl).

| Feature | Landrace_v1 | Sscrofa11.1 |
|---|---|---|
| biological_region | 162024 | 150757 |
| C_gene_segment | 18 | 1 |
| CDS | 554204 | 501906 |
| D_gene_segment | 2 | 0 |
| exon | 648128 | 556331 |
| five_prime_UTR | 61342 | 57623 |
| gene | 19252 | 20983 |
| J_gene_segment | 15 | 4 |
| lnc_RNA | 32637 | 11548 |
| miRNA | 600 | 370 |
| mRNA | 47628 | 44425 |
| ncRNA | 0 | 16 |
| ncRNA_gene | 28876 | 12899 |
| pseudogene | 440 | 416 |
| pseudogenic_transcript | 440 | 416 |
| region | 21 | 21 |
| rRNA | 48 | 17 |
| scRNA | 25 | 25 |

| | | |
|---|---|---|
| snoRNA | 768 | 556 |
| snRNA | 1124 | 1077 |
| three_prime_UTR | 45663 | 43723 |
| transcript | 17 | 0 |
| tRNA | 22 | 22 |
| V_gene_segment | 66 | 13 |
| Y_RNA | 8 | 8 |

Supplementary Table 6: GO enrichment analysis with PANTHER, displaying the most significant terms.

| GO biological process complete | FDR |
|---|---|
| detection of chemical stimulus involved in sensory perception of smell (GO:0050911) | 2.26E-227 |
| sensory perception of smell (GO:0007608) | 4.79E-226 |
| detection of chemical stimulus involved in sensory perception (GO:0050907) | 1.04E-224 |
| detection of chemical stimulus (GO:0009593) | 4.59E-223 |
| detection of stimulus involved in sensory perception (GO:0050906) | 8.81E-221 |
| sensory perception of chemical stimulus (GO:0007606) | 5.87E-220 |
| detection of stimulus (GO:0051606) | 8.77E-216 |
| sensory perception (GO:0007600) | 3.25E-200 |
| nervous system process (GO:0050877) | 7.85E-180 |
| G protein-coupled receptor signaling pathway (GO:0007186) | 3.05E-178 |
| system process (GO:0003008) | 2.29E-160 |
| response to chemical (GO:0042221) | 4.67E-123 |
| signal transduction (GO:0007165) | 9.68E-96 |
| signaling (GO:0023052) | 6.34E-90 |
| cell communication (GO:0007154) | 8.91E-90 |
| multicellular organismal process (GO:0032501) | 5.54E-82 |
| cellular response to stimulus (GO:0051716) | 1.09E-80 |
| response to stimulus (GO:0050896) | 1.18E-66 |
| regulation of cellular process (GO:0050794) | 3.80E-42 |
| regulation of biological process (GO:0050789) | 5.71E-38 |
| biological regulation (GO:0065007) | 6.21E-38 |
| cellular process (GO:0009987) | 9.51E-26 |
| biological_process (GO:0008150) | 1.90E-15 |

Supplementary Table 7: Estimated physical length and position of the 20 centromeres in Landrace_v1. The assembly of Ssc16 does not read all the way through the centromere so the actual length is longer than 1.5Mb.

| Chromosome | Centromere Start-End (Landrace_v1) |
|---|---|
| Ssc01 | 92,720,000-100,549,400 (7.8) |
| Ssc02 | 53,760,000-57,180,000 (3.4) |
| Ssc03 | 46,680,000-49,069,951 (2.4) |

| | |
|---|---|
| Ssc04 | 46,440,000-49,840,350 (3.4) |
| Ssc05 | 42,480,000-44,488,072 (2.0) |
| Ssc06 | 40,020,000-42,500,000 (2.5) |
| Ssc07 | 29,180,000-29,760,000 (0.6) |
| Ssc08 | 55,320,000-57,153,428 (1.8) |
| Ssc09 | 63,420,000-66,509,288 (3.1) |
| Ssc10 | 34,800,000-37,640,000 (2.2) |
| Ssc11 | 36,100,000-39,060,000 (3.0) |
| Ssc12 | 29,780,000-33,440,000 (3.7) |
| Ssc13 | 2,420,000-13,720,000 (11.3) |
| Ssc14 | 2,220,000-5,720,000 (3.5) |
| Ssc15 | 1,980,000-13,780,000 (11.8) |
| Ssc16 | 0-1,540,000 (>1.5)* |
| Ssc17 | 2,100,000-6,380,000 (4.3) |
| Ssc18 | 1,940,000-7,440,000 (5.5) |
| SscX | 53,140,000-57,700,000 (4.6) |
| SscY | 21,580,000-25,780,000 (4.2) |

Supplementary Table 8: The sequence regions at the beginning of the acrocentric chromosomes

| | Telomere repeat | MC1 satellite | TTCTA-repeat | TAGAA-repeat | AC2 satellite |
|---|---|---|---|---|---|
| Ssc13 | 1-1518 | 8420-712821 | 712818-1272104 | 1273727-2392474 | 2432142-13703573 |
| Ssc14 | 3-3602 | 11191-684416 | 684419-956563 | 958688-2188049 | 2227719-5704129 |
| Ssc15 | 2-6384 | 13953-815448 | 815449-978726 | 981044-1949310 | 1993371-13774139 |
| Ssc16 | - | - | - | - | 224-1538555 |
| Ssc17 | 3-3390 | 10974-784360 | 784361-933503 | 935154-2064438 | 2108697-6366927 |
| Ssc18 | | 1-588275 | 588276-1009346 | 1010629-1903348 | 1947478-7421339 |

#### Supplementary figures

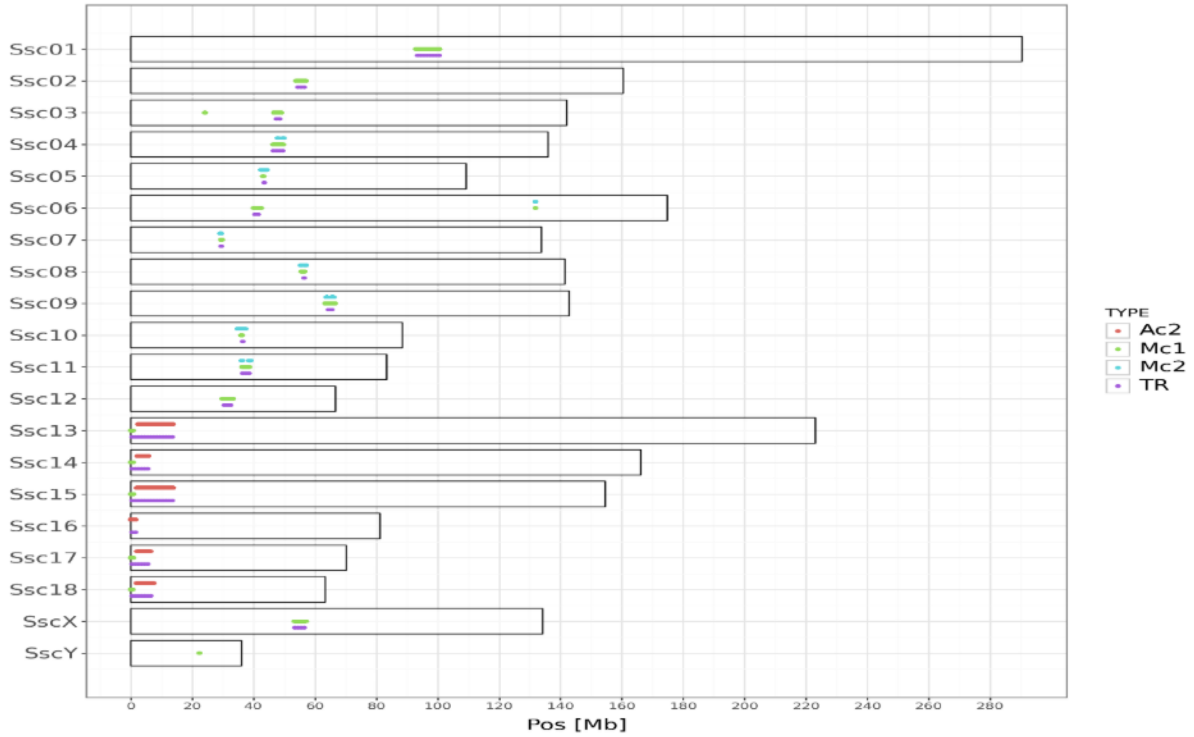


Supplementary Figure 1: Suggested centromere locations based on the two strategies used in this paper. Ac2, Mc1 and Mc2 show windows of 10kb where more than 10% of the sequence maps to one of the centromere specific sequences. TR are 1 Mb windows where more than 50% of the sequence is reported as being part of a repeat. Each dot shows a window of 10kb with >10% of the bases overlapping an alignment from the downloaded sequences.

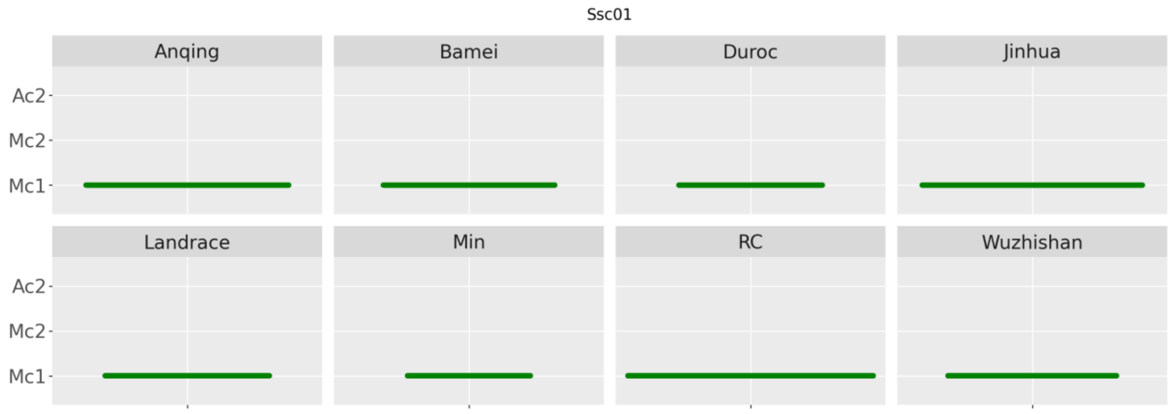

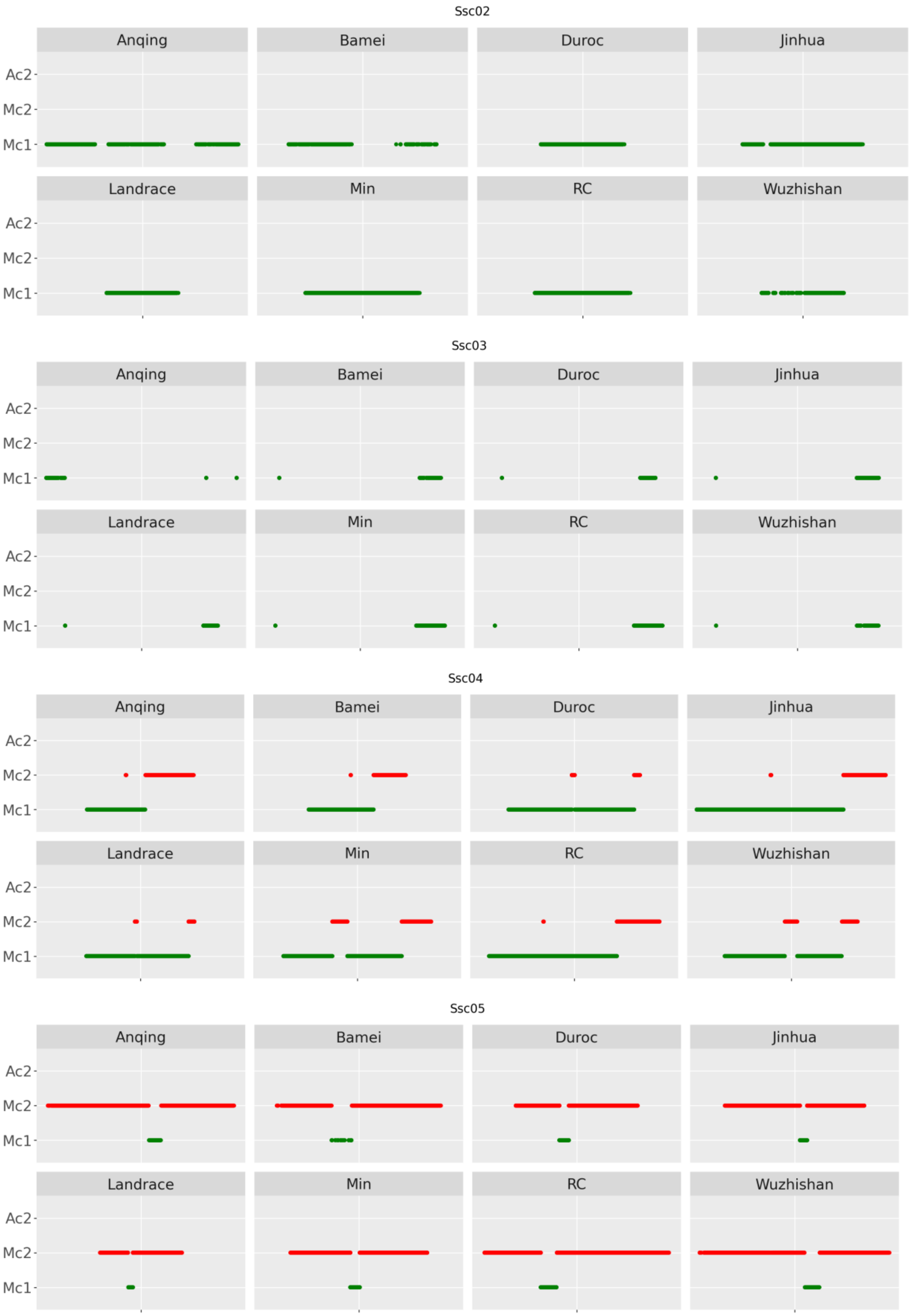
Ssc02
Anqing
Bamei
Duroc
Jinhua
Landrace
Min
RC
Wuzhishan
Ac2
Mc2
Mc1
Ssc03
Ssc04
Ssc05

Ssc06

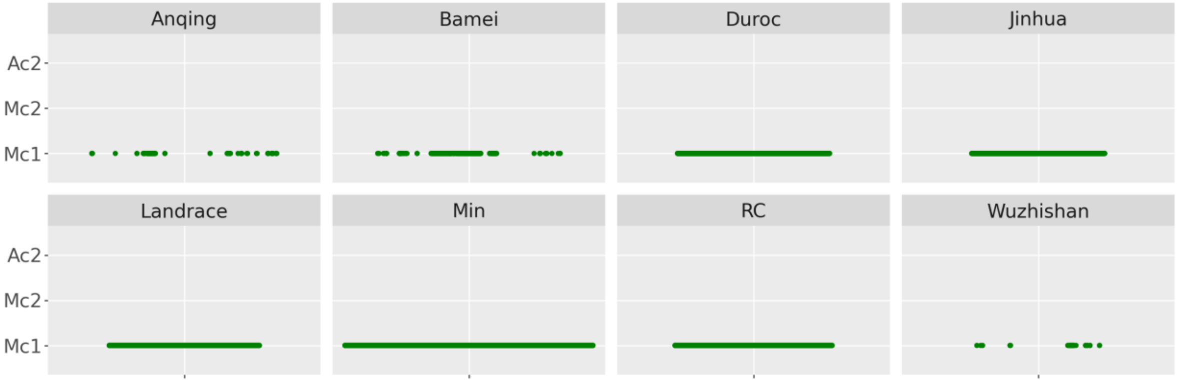


Ssc07

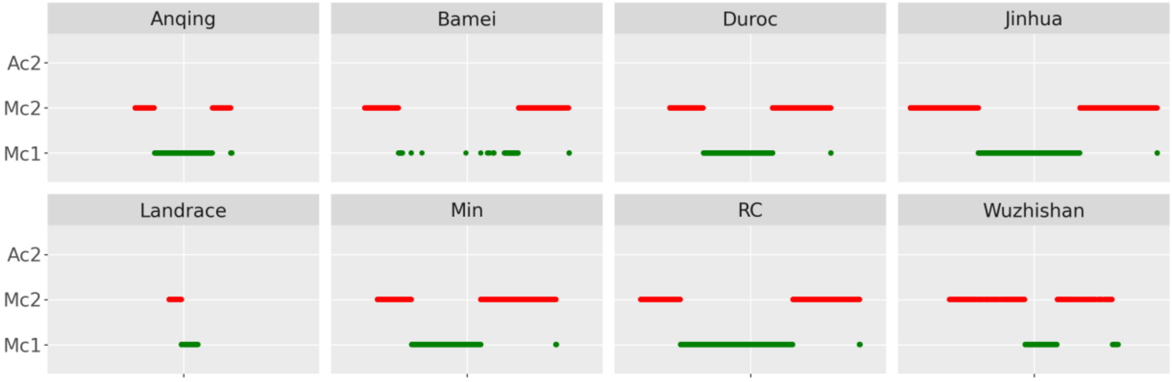


Ssc08

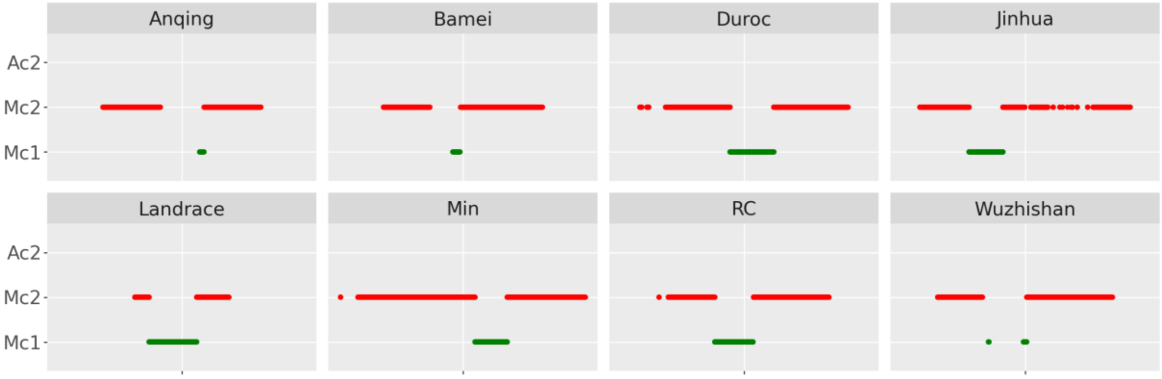


Ssc09

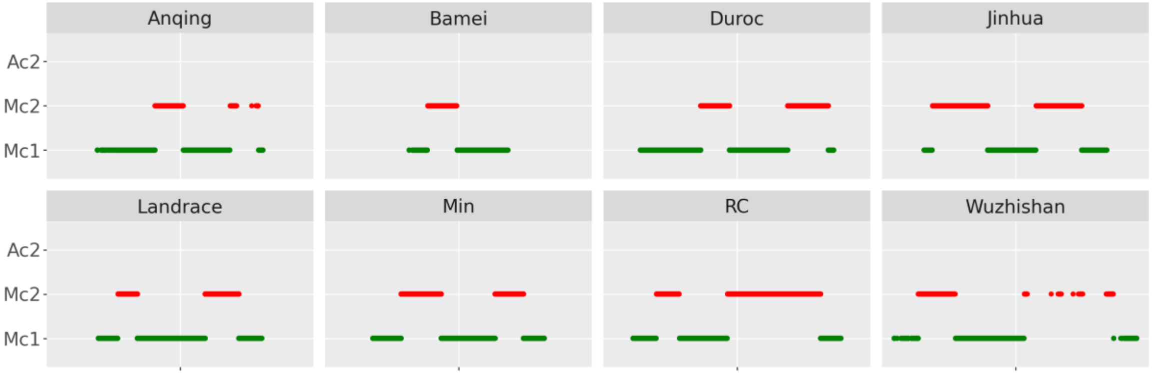

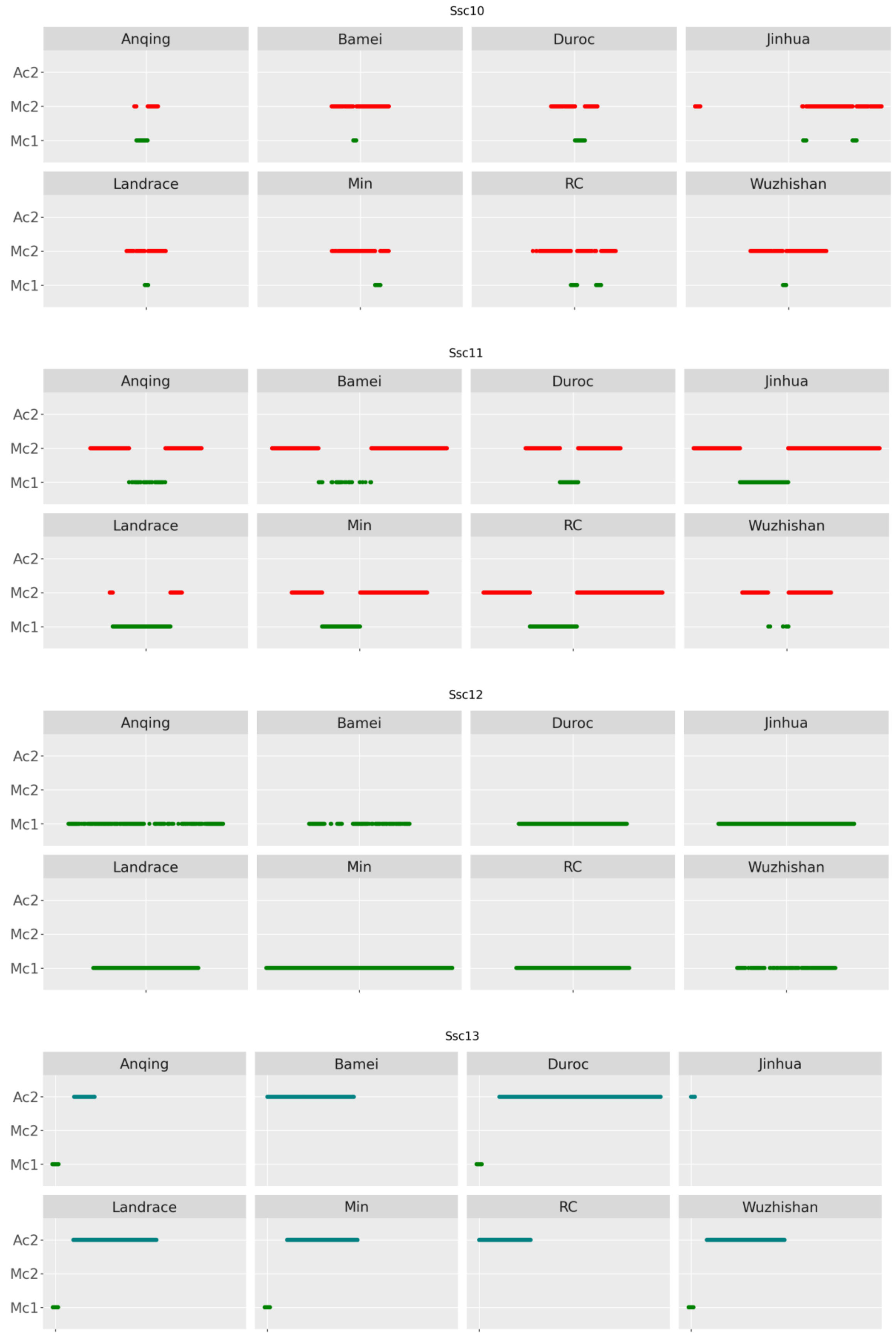
Ssc10
Anqing
Bamei
Duroc
Jinhua
Landrace
Min
RC
Wuzhishan
Ac2
Mc2
Mc1
Ssc11
Anqing
Bamei
Duroc
Jinhua
Landrace
Min
RC
Wuzhishan
Ac2
Mc2
Mc1
Ssc12
Anqing
Bamei
Duroc
Jinhua
Landrace
Min
RC
Wuzhishan
Ac2
Mc2
Mc1
Ssc13
Anqing
Bamei
Duroc
Jinhua
Landrace
Min
RC
Wuzhishan
Ac2
Mc2
Mc1

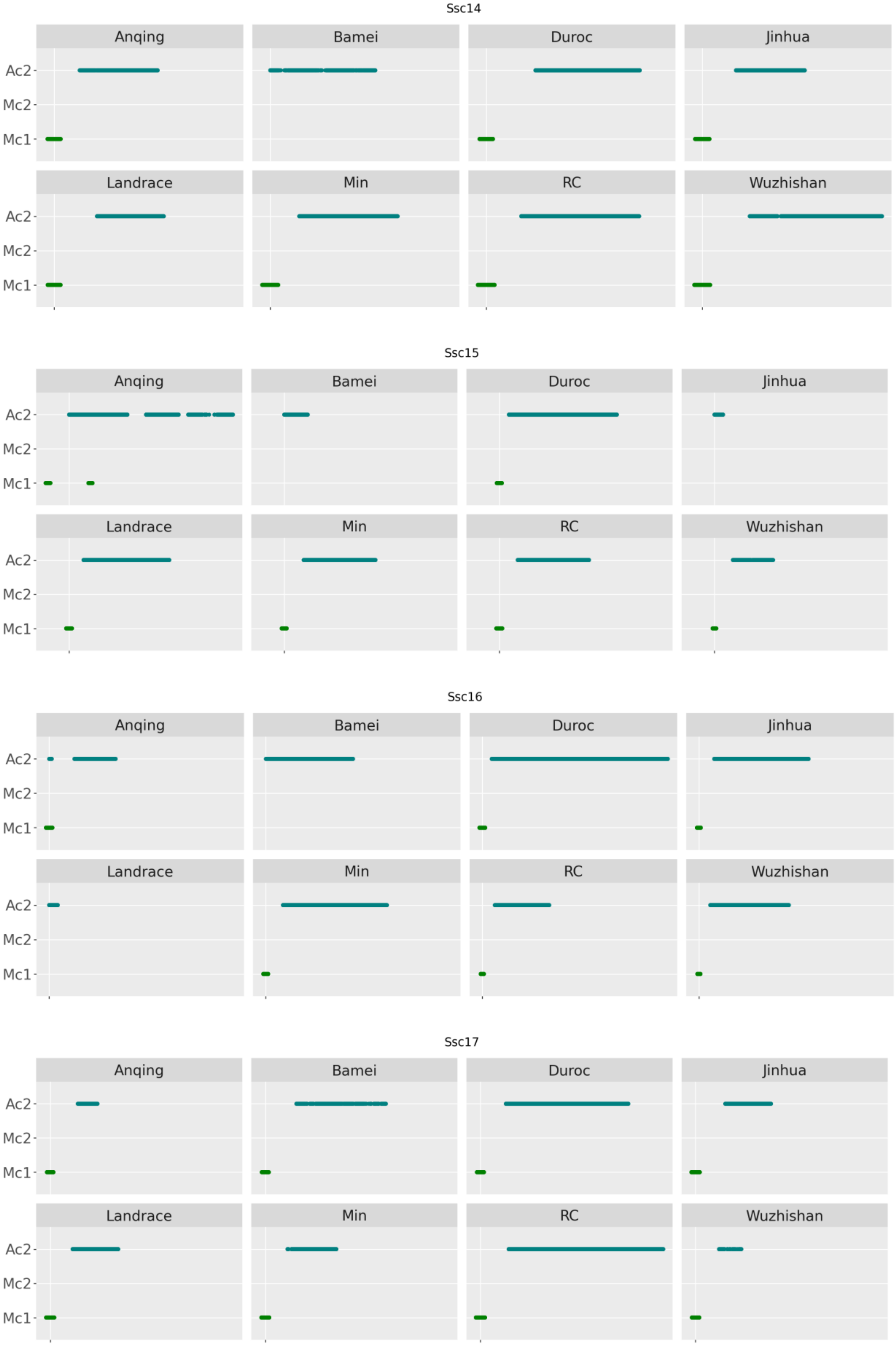

Ssc14
Anqing
Bamei
Duroc
Jinhua
Landrace
Min
RC
Wuzhishan
Ac2
Mc2
Mc1
Ssc15
Anqing
Bamei
Duroc
Jinhua
Landrace
Min
RC
Wuzhishan
Ac2
Mc2
Mc1
Ssc16
Anqing
Bamei
Duroc
Jinhua
Landrace
Min
RC
Wuzhishan
Ac2
Mc2
Mc1
Ssc17
Anqing
Bamei
Duroc
Jinhua
Landrace
Min
RC
Wuzhishan
Ac2
Mc2
Mc1

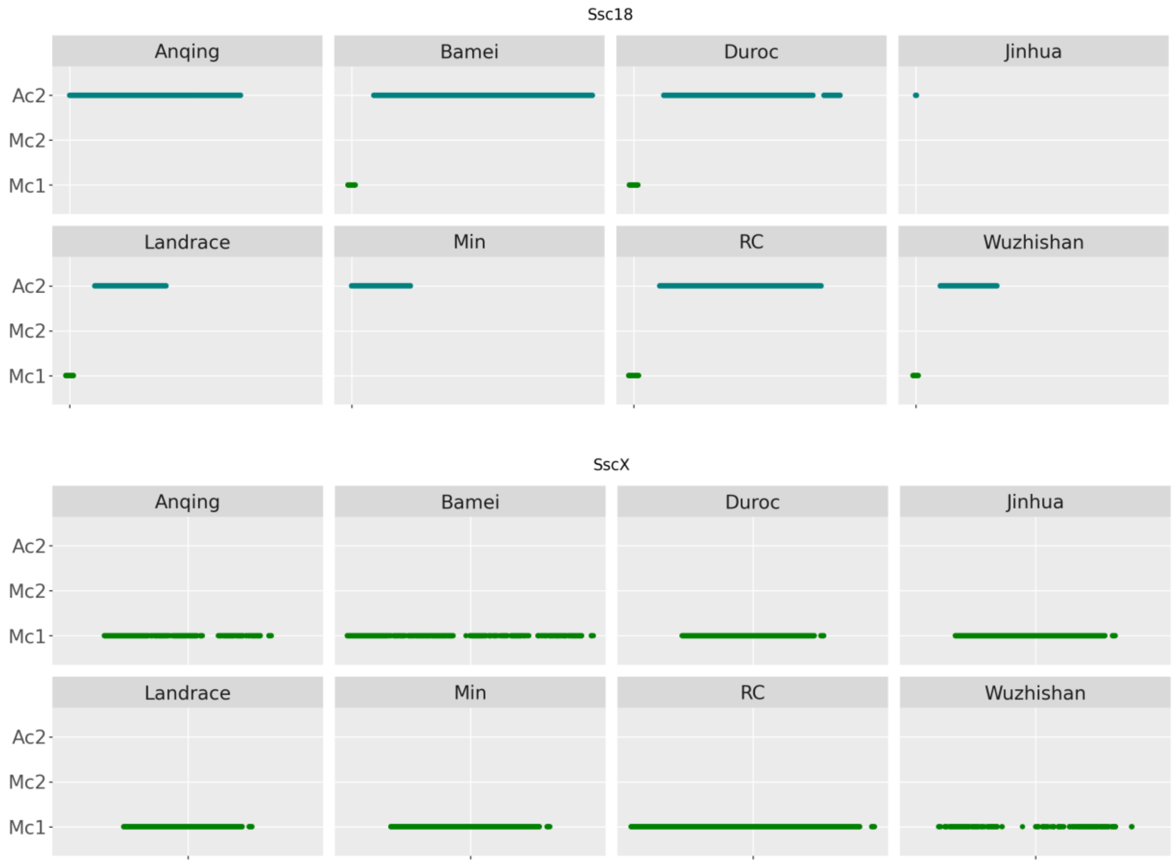


Supplementary Figure 2: The repeat structure of the centromere for each of the 18 autosomes and the X-chromosome as described by the location and frequency of the Mc1, Mc2 and Ac2 repeats published at NCBI.

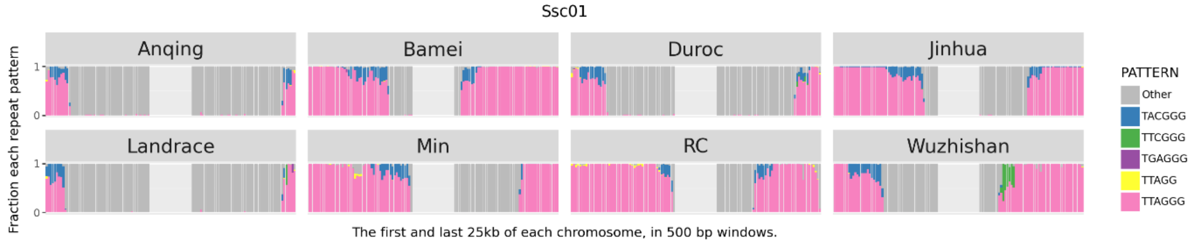


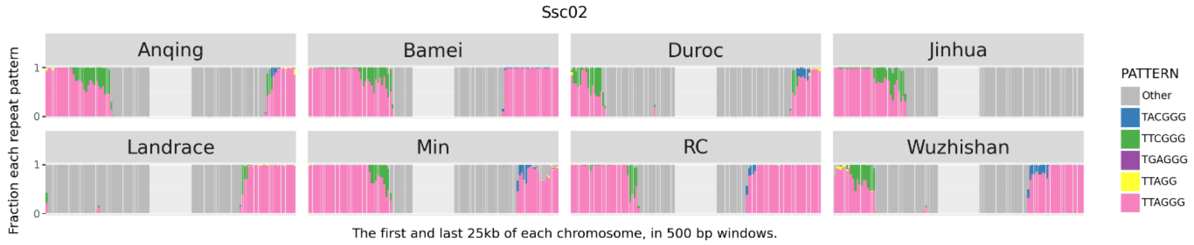

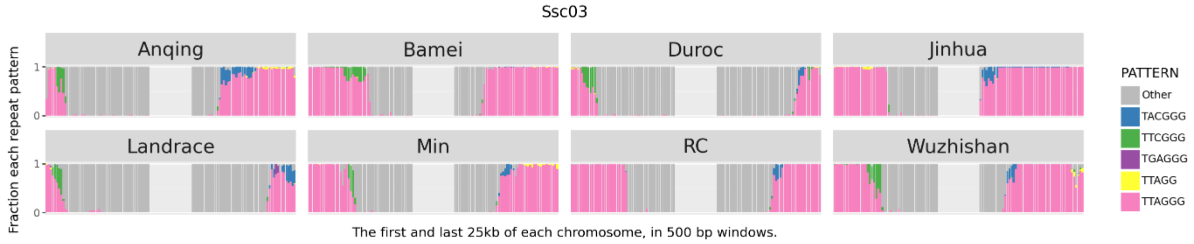
Ssc03
Anqing
Bamei
Duroc
Jinhua
Landrace
Min
RC
Wuzhishan
Fraction each repeat pattern
1
0
The first and last 25kb of each chromosome, in 500 bp windows.
PATTERN
Other
TACGGG
TTCGGG
TGAGGG
TTAGG
TTAGGG

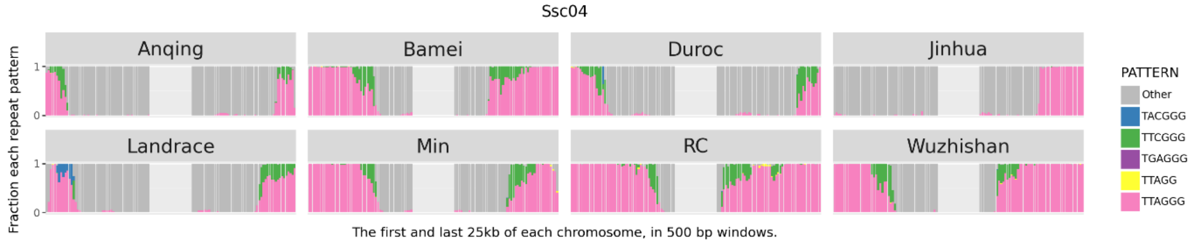
Ssc04
Anqing
Bamei
Duroc
Jinhua
Landrace
Min
RC
Wuzhishan
Fraction each repeat pattern
1
0
The first and last 25kb of each chromosome, in 500 bp windows.
PATTERN
Other
TACGGG
TTCGGG
TGAGGG
TTAGG
TTAGGG

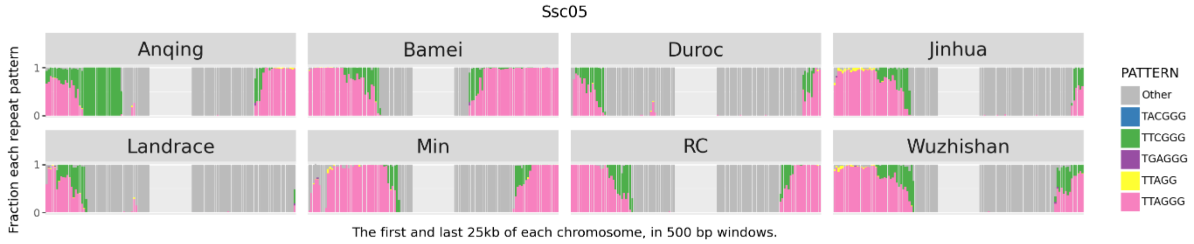
Ssc05
Anqing
Bamei
Duroc
Jinhua
Landrace
Min
RC
Wuzhishan
Fraction each repeat pattern
1
0
The first and last 25kb of each chromosome, in 500 bp windows.
PATTERN
Other
TACGGG
TTCGGG
TGAGGG
TTAGG
TTAGGG

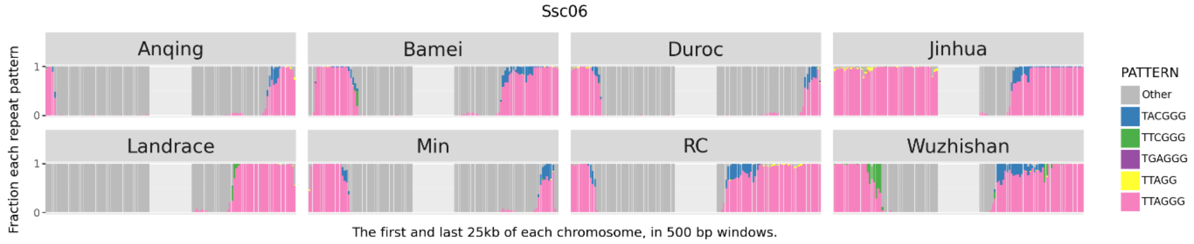
Ssc06
Anqing
Bamei
Duroc
Jinhua
Landrace
Min
RC
Wuzhishan
Fraction each repeat pattern
1
0
The first and last 25kb of each chromosome, in 500 bp windows.
PATTERN
Other
TACGGG
TTCGGG
TGAGGG
TTAGG
TTAGGG

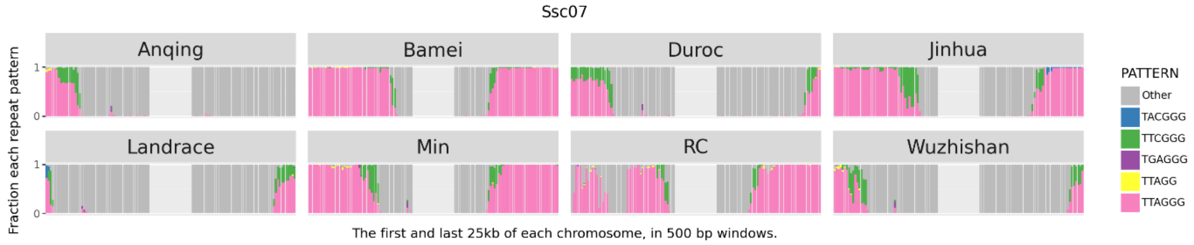
Ssc07
Anqing
Bamei
Duroc
Jinhua
Landrace
Min
RC
Wuzhishan
Fraction each repeat pattern
1
0
The first and last 25kb of each chromosome, in 500 bp windows.
PATTERN
Other
TACGGG
TTCGGG
TGAGGG
TTAGG
TTAGGG

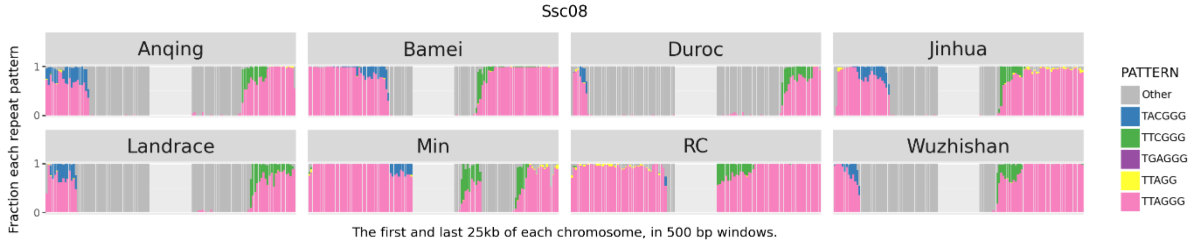
Ssc08
Anqing
Bamei
Duroc
Jinhua
Landrace
Min
RC
Wuzhishan
Fraction each repeat pattern
1
0
The first and last 25kb of each chromosome, in 500 bp windows.
PATTERN
Other
TACGGG
TTCGGG
TGAGGG
TTAGG
TTAGGG

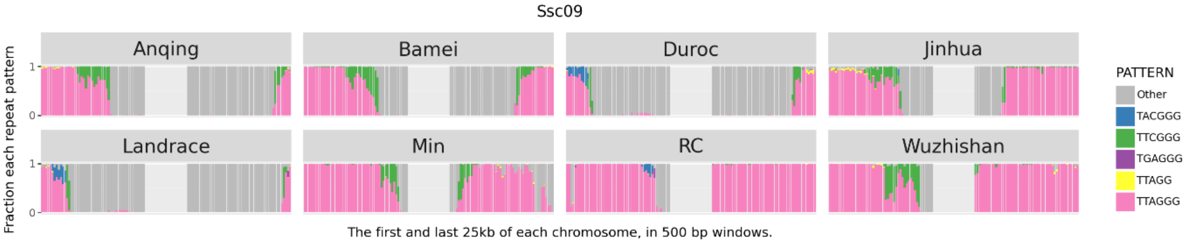
Ssc09
Anqing
Bamei
Duroc
Jinhua
Landrace
Min
RC
Wuzhishan
Fraction each repeat pattern
1
0
The first and last 25kb of each chromosome, in 500 bp windows.
PATTERN
Other
TACGGG
TTCGGG
TGAGGG
TTAGG
TTAGGG

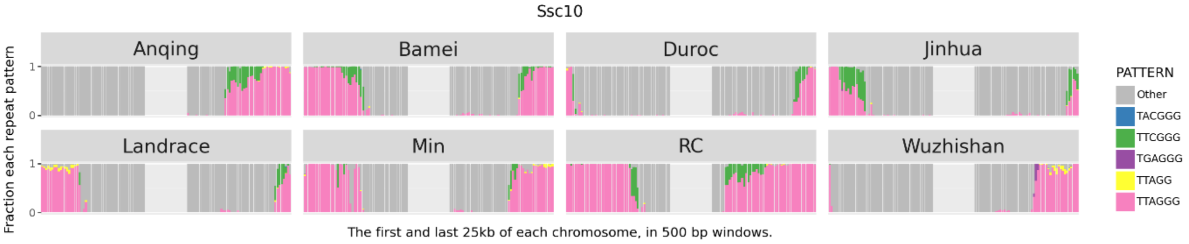
Ssc10
Anqing
Bamei
Duroc
Jinhua
Landrace
Min
RC
Wuzhishan
Fraction each repeat pattern
1
0
The first and last 25kb of each chromosome, in 500 bp windows.
PATTERN
Other
TACGGG
TTCGGG
TGAGGG
TTAGG
TTAGGG

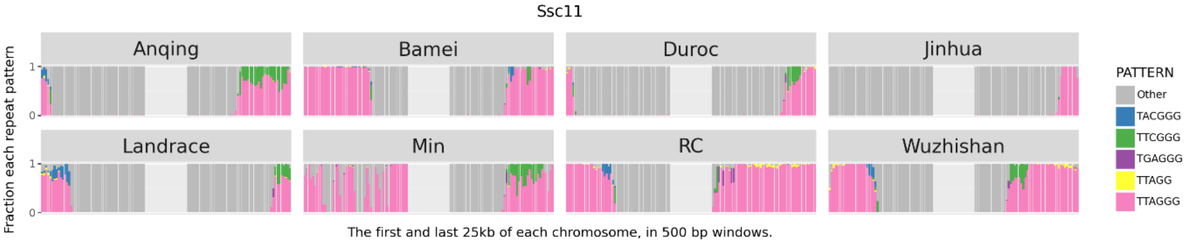
Ssc11
Anqing
Bamei
Duroc
Jinhua
Landrace
Min
RC
Wuzhishan
Fraction each repeat pattern
1
0
The first and last 25kb of each chromosome, in 500 bp windows.
PATTERN
Other
TACGGG
TTCGGG
TGAGGG
TTAGG
TTAGGG

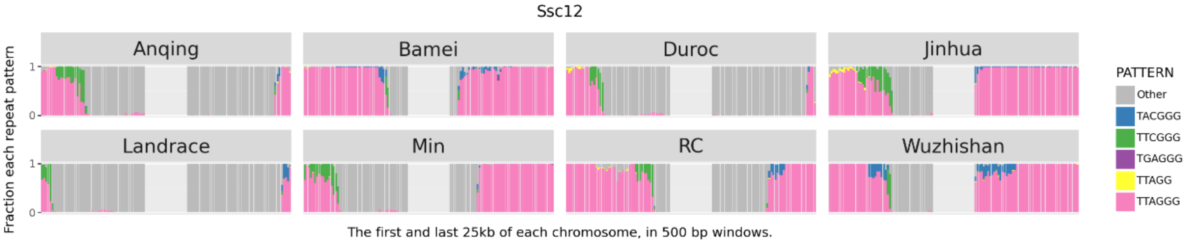
Ssc12
Anqing
Bamei
Duroc
Jinhua
Landrace
Min
RC
Wuzhishan
Fraction each repeat pattern
1
0
The first and last 25kb of each chromosome, in 500 bp windows.
PATTERN
Other
TACGGG
TTCGGG
TGAGGG
TTAGG
TTAGGG

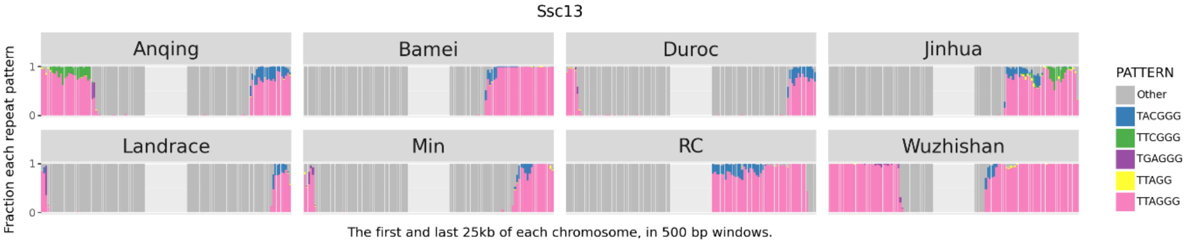
Ssc13
Anqing
Bamei
Duroc
Jinhua
Landrace
Min
RC
Wuzhishan
Fraction each repeat pattern
1
0
The first and last 25kb of each chromosome, in 500 bp windows.
PATTERN
Other
TACGGG
TTCGGG
TGAGGG
TTAGG
TTAGGG

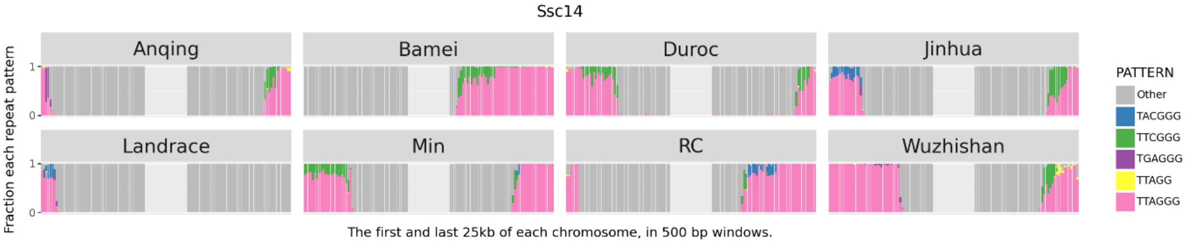
Ssc14
Anqing
Bamei
Duroc
Jinhua
Landrace
Min
RC
Wuzhishan
Fraction each repeat pattern
1
0
The first and last 25kb of each chromosome, in 500 bp windows.
PATTERN
Other
TACGGG
TTCGGG
TGAGGG
TTAGG
TTAGGG

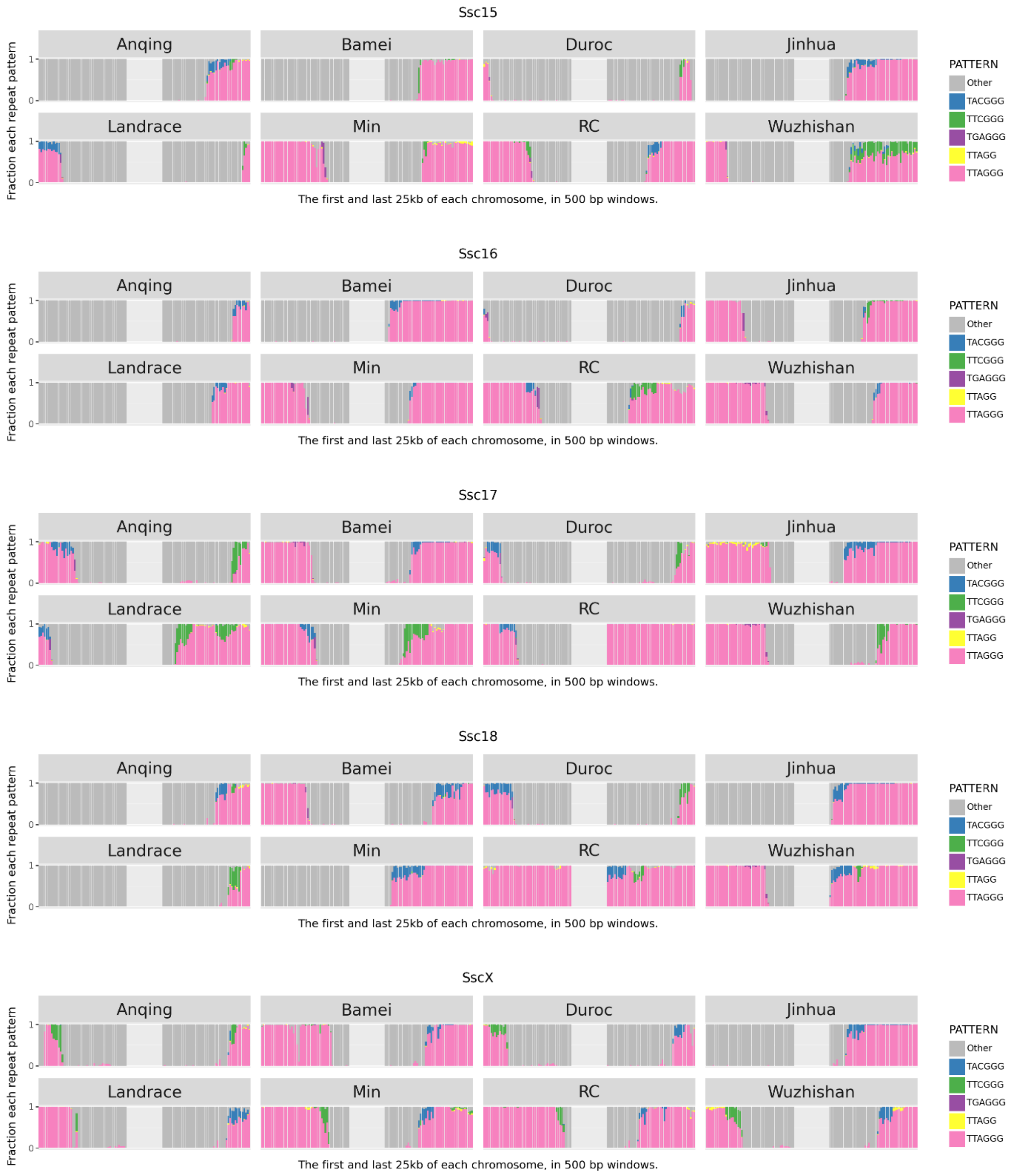


Supplementary Figure 3: Telomere region for all chromosomes within 8 pig breeds. All the breeds show the main TTAGGG repeat but also the presence of another repeat in the region where the telomere repeat transitions to the main sequence. The length of the telomere repeat is also variable.

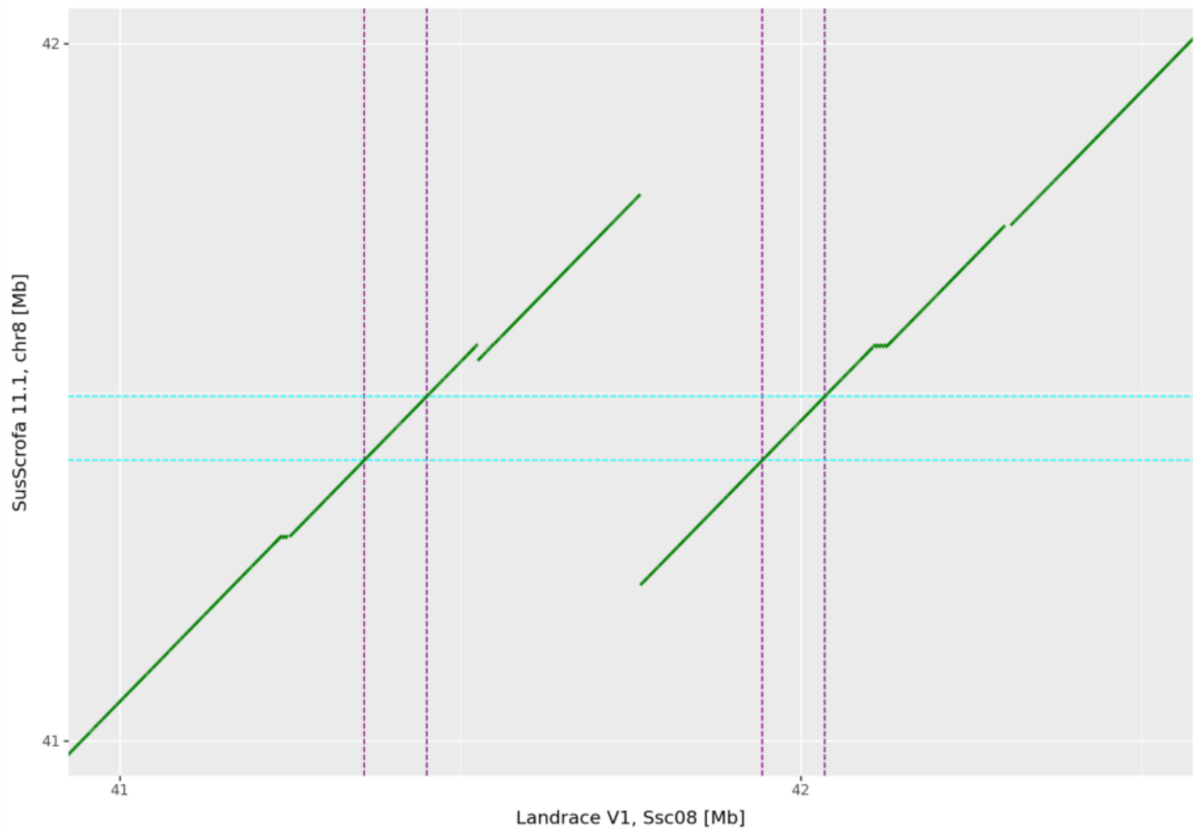


Supplementary Figure 4: Dotplot showing the local alignment of the region covering the KIT gene between Lv1 assembly and the Sus Scrofa 11.1 reference. The single copy of the KIT gene is shown in cyan, while the two copies in Landrace are shown in purple. It is also possible to observe the other duplications internal to the main duplication.
